# Personalized Medicine for Meningiomas: Drug Screening on Tumor Organoids Exposes Therapeutic Vulnerabilities to HDAC1/2i Panobinostat

**DOI:** 10.1101/2024.11.26.625347

**Authors:** Gerhard Jungwirth, Junguo Cao, Catharina Lotsch, Rolf Warta, Mahmoud Moustafa, Maximilian Knoll, Tao Yu, Viktor Braun, Lena Jassowicz, Philip Dao Trong, Alexander Younsi, Moritz Scherer, Martin Bendszus, Sandro M. Krieg, Andreas von Deimling, Juergen Debus, Felix Sahm, Amir Abdollahi, Andreas Unterberg, Christel Herold-Mende

**Author notes:** These authors contributed equally to this work (Gerhard Jungwirth, Junguo Cao). **Address correspondence to:** Prof. Dr. Christel Herold-Mende, Division of Experimental Neurosurgery, Department of Neurosurgery, Heidelberg University, INF400, 69120 Heidelberg, Germany.

## Abstract

Managing aggressive meningiomas remains challenging due to limited treatment options besides surgical tumor removal and radiotherapy. To identify novel therapies for aggressive meningiomas, we established a multi-step drug screening workflow, focusing on targetable genes obtained from transcriptome data of highly aggressive grade 3 meningiomas. *In vitro* screening of 107 targeted drugs identified nine effective inhibitors. To study these drugs in a more natural environment, we established a standardized patient-derived tumor organoid (TO) model preserving accurately the original tissue’s genotype and phenotype. Individual drug responses were assessed in TOs from 60 molecularly characterized meningioma cases. Especially the FDA-approved epigenetic drug panobinostat demonstrated high antimeningioma efficacy in 70% of TOs, mediated through HDAC1/2 inhibition. In addition, treatment in an orthotopic *in vivo* model revealed a significantly improved survival. In a heavily pretreated patient suffering from an anaplastic meningioma, oral panobinostat treatment could delay the tumor growth rate. In search of the molecular mechanism underlying a potential intrinsic panobinostat resistance, we identified upregulation of the HDAC8-TGFβ-EMT axis in the TO model and subsequent HDAC8 depletion substantially increased the sensitivity to panobinostat. These data highlight the utility of personalized drug screenings on TOs to identify suitable drug targets and inhibitors for a more effective treatment of clinically aggressive meningiomas and help to advance our understanding of counteracting resistance mechanisms.

**One Sentence Summary:** This study provides strong *in vitro*, *in vivo*, *ex vivo*, and patient evidence for the efficacy of the HDACi panobinostat to treat clinically aggressive meningiomas and uncovered a potential intrinsic resistance mechanism by activation of the HDAC8-TGFβ-EMT axis.

## INTRODUCTION

Meningiomas are the most common primary brain tumor^1^. Therapy options include maximal-safe surgical resection and additional radiotherapy, especially for high-grade meningiomas^2^. Still, recurrence of atypical and anaplastic meningiomas is common, translating into a median overall survival of only four years in the latter case^3^. To date, no successful systemic treatment exists^4^. Over 20 different drugs have been clinically tested^4^, mostly in small and non-randomized trials, including hydroxyurea^5,6^, temozolomide^7^, tamoxifen^8^, trabectedin^9^, and tyrosine kinase inhibitors^10–12^, failing to provide any significant clinical benefit. The only double-blind randomized phase III trial conducted in meningioma with the antiprogestin agent mifepristone was negative^13^.

To test the efficacy of novel drugs, researchers primarily relied on cell lines as a robust and inexpensive tool for cancer research^14–16^. By using meningioma cell lines, we recently published the first large-scale drug screening providing a detailed overview of the antitumor activity of antineoplastic FDA-approved drugs^17^. Thereby, we identified the drugs romidepsin, omacetaxine, carfilzomib, and ixabepilone demonstrating the most impressive antimeningioma effects *in vitro* and *in vivo*^17^. However, important limitations of cell lines are that only a few meningioma cell lines exist and they cannot fully recapitulate a tumor entity because of the lack of intra- and intertumoral heterogeneity, tumor microenvironment, and a proper 3D tumor architecture^16,18^. Tumor organoids (TOs) or patient-derived organoids (PDOs) are novel, complex three-dimensional *ex vivo* tissue culture models that under optimal conditions accurately reflect the genotype and phenotype of the original tissue while preserving its cellular heterogeneity^18^. They offer a new and exciting platform for studying cancer biology and guiding personalized therapies^19^. However, published protocols to create PDOs vary widely^18,20–22^. Most commonly, tumor tissue is minced into small pieces, usually followed by embedding into a synthetic extracellular matrix^18,22^. However, results from heterogenous tumor fragments may hinder standardized drug screenings^20,23^. Furthermore, to attain required numbers of PDOs for drug screening, culturing for weeks to months might be necessary aggravating shifts in the cellular composition of PDOs which also may alter drug responses^18,22,24^. All these factors limit the suitability of published protocols to generate PDOs for high-throughput drug screenings.

Here, we first conducted a large-scale drug screening on meningioma cell lines using an anticancer drug library of 107 drugs specifically selected for targets identified in aggressive meningiomas. To further interrogate anti-tumor drug responses in a more natural environment and thus under the influence of the surrounding tumor stroma, we employed a novel, standardized meningioma tumor organoid model which replicates the tissue of origin regarding its transcriptional and mutational profile, and its tumor microenvironment. Out of 107 targeted drugs, the HDACi panobinostat exerted strong antimeningioma effects *in vitro*, *in vivo* and *ex vivo* which could be attributed to the inhibition of HDAC1/2. Lastly, we used the same tumor organoid model to dissect a potential intrinsic resistance mechanism to panobinostat and identified an important role of HDAC8 inducing a TGFβ-mediated epithelial-to-mesenchymal transition. Overall, we were able to show that the HDACi panobinostat is very effective in treating clinically aggressive meningiomas *in vitro*, *in vivo* and *ex vivo*.

## RESULTS

### *In silico* exploration for druggable targets and drug screening identifies nine most effective drugs

To identify drug targets for aggressive meningiomas, we assessed our previously published transcriptome dataset (GSE74385)^25^ enriched with an extraordinarily high proportion of advanced grade 3 MGMs (n=28) to screen for expressed genes (n=442) that can be targeted by available inhibitors (Fig. 1A). Selection of drugs targeting these genes was based on two criteria: Only drugs were considered which were clinically evaluated, and for each target, a maximum of three compounds was selected based on available data from clinical trials. This filtering process resulted in 107 drugs targeting 46 genes (Suppl. Table S1). Most compounds were studied in tumors (oncology, *n*=94, 88%) while the remaining compounds were non-oncology agents (*n*=13, 12%; Suppl. Fig. 1A). Furthermore, 81 of these drugs are currently studied in clinical trials (76%), while 26 drugs have already been approved by the FDA (24%, Suppl. Fig. 1B).

**Fig. 1:**
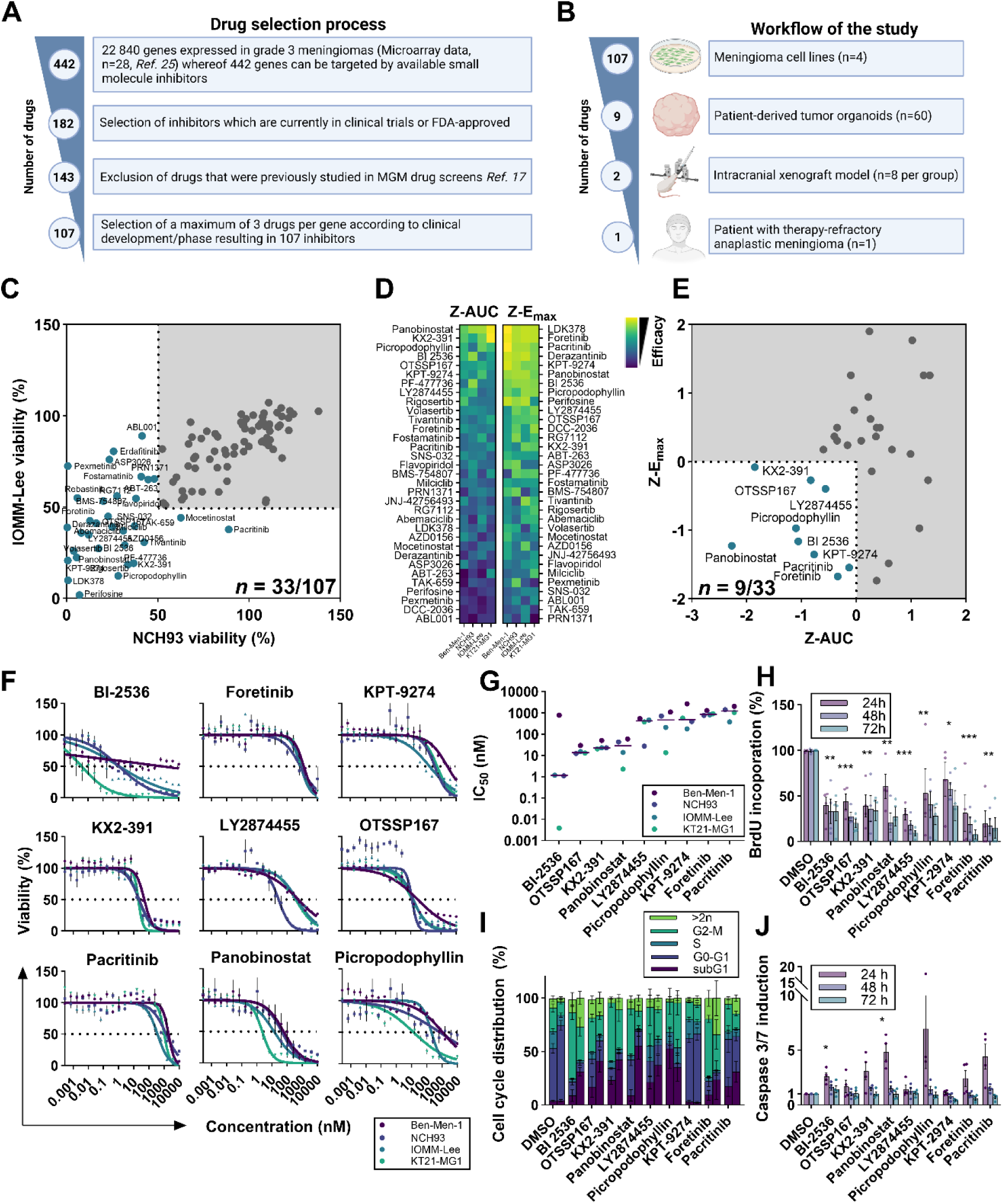
Screening of a panel of targeted drugs identified nine highly potent compounds. **(A)** Drug selection workflow. **(B)** Drug screening workflow. **(C)** Anti-meningioma activity of the drug library of targeted drugs (*n*=107) was screened at a single dose (2.5 µM) in two anaplastic meningioma cell lines IOMM-Lee and NCH93. Thirty-three drugs decreased viability at least by 50%. **(D)** Dose-response curves of 33 drugs studied in four meningioma cell lines (Ben-Men-1, NCH93, IOMM-Lee, and KT21-MG1) in half-logarithmic concentrations ranging from 0.1 nM to 10 µM. Heatmap represents the z-transformed area under the curve (z-AUC) and maximum inhibitory effect (z-E_max_). **(E)** Nine drugs demonstrated a combined z-AUC and z-E_max_ below the mean (0). **(F)** Dose-response curves were validated using extended dose ranges and crystal violet staining. **(G)** IC_50_ values of each drug and cell line (dots), and their corresponding medians are plotted. **(H)** Meningioma cell lines (*n*=4) were treated with the respective drug at 10x IC_50_ and a BrdU assay was performed after 24, 48, and 72 h. Data were combined from four meningioma cell lines and results were normalized to DMSO-treated cells. **(I)** Quantification of cell cycle perturbations from drug treatment of four cell lines treated at 10x IC_50_ for 24 (left) and 48 h (right bar). **(J)** Induction of apoptosis was measured after treating meningioma cells at 10x IC_50_ by Caspase-Glo 3/7 assay for 24, 48, and 72 h. Data are presented in mean±SEM or otherwise specified. One-way ANOVA followed by a *post hoc* Dunnett’s multiple comparison test was performed for F, G, and H. *, *P* < 0.05; **, *P* < 0.01; ***, *P* < 0.001.

The drug screening workflow of the study is presented in Fig. 1B. First, this panel of targeted drugs was screened on meningioma cell lines in a two-step process. In step one, drugs were tested at a single dose (2.5 µM) for antineoplastic effects on two malignant meningioma cell lines (NCH93 and IOMM-Lee). 33/107 drugs reduced cell viability by 50% or more in each cell line (Fig. 1C). These drugs (*n*=33) were further analyzed in a six-point dose-response scheme in four meningioma cell lines (Ben-Men-1, NCH93, IOMM-Lee, KT21-MG1; Fig. 1D). The most effective drugs were selected based on the lowest mean after respective z-transformation of the area under the curve (z-AUC) and the maximum inhibitory effect (z-E_max_) of all cell lines. Screening resulted in the identification of the following drugs: OTSSP167 (MELKi), panobinostat (pan-HDACi), picropodophyllin (IGF-1Ri), KPT-9274 (PAK4i and NAMPTi), pacritinib (JAKi, FLT3i), BI-2536 (PLK1i and BRDi), foretinib (METi and other RTKs), LY2874455 (Pan-FGFRi), and KX2-391 (SRCi) (Fig. 1E; Suppl. Fig. 1C). These candidates exclusively belonged to the group of oncological drugs, whereof panobinostat and pacritinib already received FDA-approval for the treatment of multiple myeloma and cancerous myelofibrosis by the time of writing^26^. Validation of the dose-response curves resulted in median IC_50_ in the lower nanomolar range across all cell lines for the top four drugs: 1.14 nM for BI-2536 (range: 0.004-774), 13.8 nM for OTSSP167 (12-29), 22 nM for KX2-391 (20-50), and 28 nM for panobinostat (2.2-58; Suppl. Table 1). The cell lines exhibited similar drug responses except for BI-2536 (Fig. 1G). Antiproliferative effects were confirmed by BrdU assay (Fig. 1H). This was accompanied by an increase of the subG1 fraction for most drugs as analyzed by flow cytometry (Fig. 1I), and the induction of caspase 3/7 especially at 24 h (Fig. 1J).

Taken together, by employing a large-scale multi-step drug screening for a considerable number of potential targets expressed in aggressive meningioma, we could identify nine promising drugs inhibiting proliferation and inducing apoptosis *in vitro*.

### Establishment of a novel, standardized patient-derived tumor organoid model

Next, we studied the efficacy of the top 9 drugs in tumor organoids (TOs). This represents a more natural environment because it contains patient-derived tumor and non-tumor cells known to be implicated in drug resistance^27^. However, most of the published protocols from other entities typically take prolonged time to establish TOs, thereby aggravating shifts in cellular compositions and leading to a variable tumor organoid size which hinders large-scale drug screenings^18,22,24^. Therefore, we developed a tumor organoid model which can be generated within days, in a standardized size and composition, and in large quantities and thus is suitable for timely large-drug screenings. This was achieved by controlled reaggregation of freshly prepared single cell suspensions of MGM tissue samples by magnetic particles in anti-adhesive microtiter plates leading to the formation of several hundred tumor organoids equal in size (Fig. 2A). TOs consisted of largely viable cells, whereas dead cells were predominantly found outside of the organoid (Fig. 2B). Single cells reaggregated to tissue-like structures within 2-3 days resulting in a reduced diameter by 67-77% independent of the cell number (Fig. 2C; Suppl. Fig. 2A&B). Thereafter, tumor organoid size remained stable throughout a 14 days observation period. Similarly, after an initial drop of intracellular ATP concentration, TO’s viability remained stable (Fig. 2D). This might be caused by the mechanical and enzymatic stress of cells, which they have undergone when being separated into a single cell suspension. Of utmost importance, transcriptomes of parental tumors and corresponding tumor organoids were highly similar and clustered together (Fig. 2E). Also, mutations detected in the parental MGM tissues were preserved in the corresponding tumor organoids (Fig. 2F). H&E staining confirmed the successful establishment of dense tissue-like structures (Fig. 2G). Moreover, preservation of the tumor microenvironment was shown by multicolor staining for meningioma cells (EMA), extracellular matrix (fibronectin), and macrophages (CD68) (Fig. 2H).

**Fig. 2:**
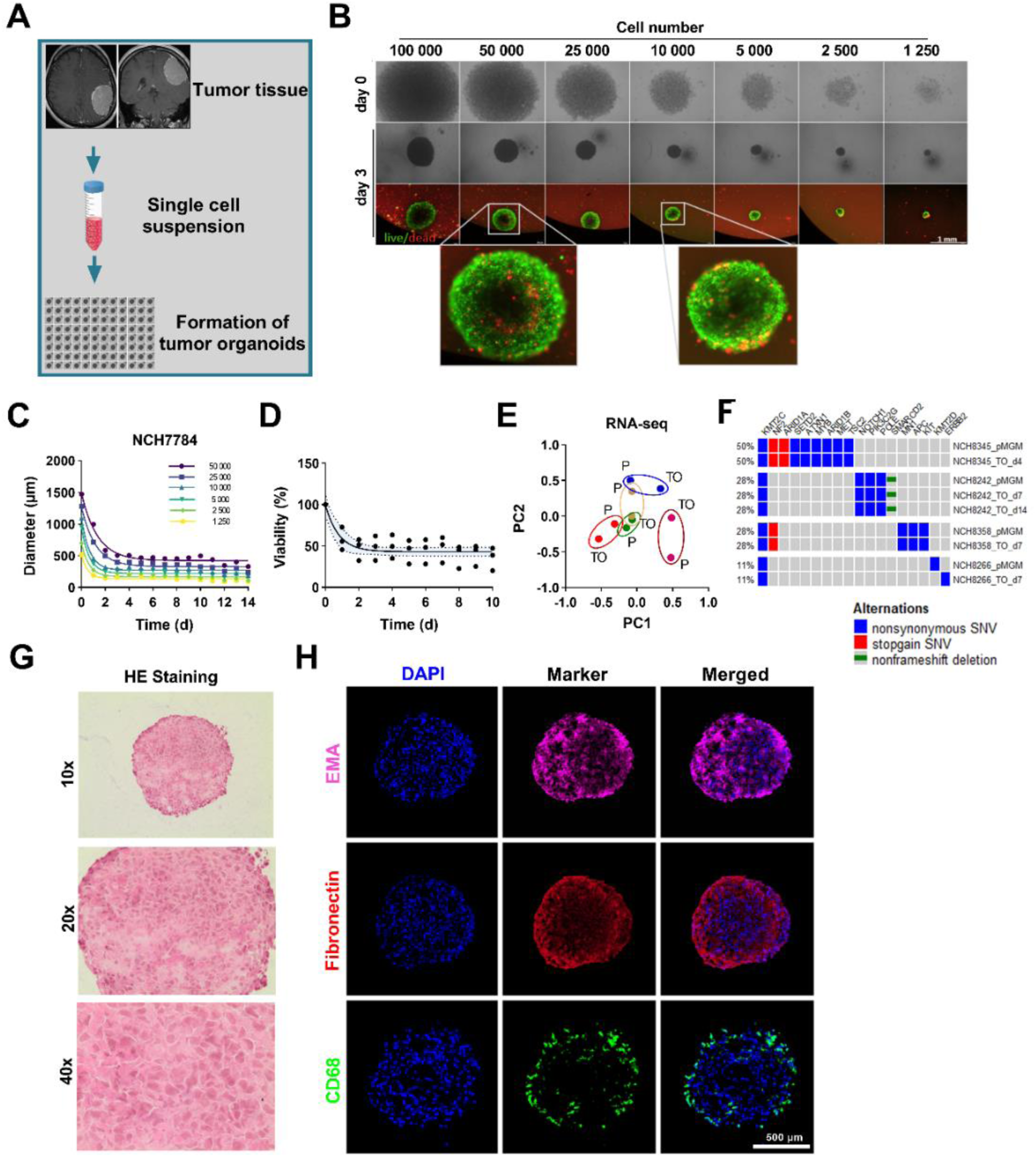
Standardized patient-derived tumor organoid model mirrors patient-tumor. **(A)** Workflow of generating meningioma tumor organoids. **(B)** Effective tumor organoid preparation from 1,250 up to 100,000 cells/organoid within 3 days. Live (green)/dead (red) staining demonstrates viability of tumor organoids. Bar represents 1 mm. **(C)** During formation tumor organoids reduce their diameter within a few days as measured by light microscope. **(D)** Viability of TOs generated from three patients was daily measured using CellTiterGlo3D and normalized to day 0. **(E)** Bulk RNA-seq of five patient-derived meningioma tissue samples and corresponding TOs was performed. Principal component analysis reveals clustering of the corresponding pairs. **(F)** Mutational profiles from meningioma tissues were preserved in TOs assessed by DNA panel sequencing. **(G)** Representative H&E staining of a TO in 10x, 20x, and 40x magnification illustrates tissue-like structures of TOs. **(H)** Immunofluorescence staining for the meningioma marker epithelial membrane antigen (EMA), the extracellular matrix protein fibronectin, and the macrophage marker CD68. Bar represents 500 µm.

Collectively, we developed a protocol to generate standardized TOs from MGMs in large quantities mirroring mutational and transcriptional features as well as the tumor microenvironment of the parental tumor.

### Drug screening in 60 patient-derived tumor organoids identifies OTSSP167 and panobinostat as highly effective drugs

Next, we assessed the reliability of TOs for robust drug testing. To this end, we treated TOs with the top nine compounds derived from the initial drug screening. First, we tested if drug responses depend on TO size. Drug responses remained comparable independent of the cell numbers per organoid (Suppl. Fig. 3A&B). To achieve a sufficient representation of rare cell types such as T-cells and macrophages an overall number of 25 000 cells/TO was selected for further experiments. Furthermore, maximum treatment efficacy was achieved after 72 h (Suppl. Fig. 3A&B), which was chosen for the following drug screening in TOs.

In total, we generated standardized TOs from 60 molecularly characterized meningiomas, including eight grade 2 and three grade 3 MGMs, and treated them with the nine respective drugs at increasing half-logarithmic concentrations from 10 nM to 30 µM (Fig. 3A). The assay performance was calculated by the overall Z-factor of 0.64, indicating high quality (0.5 – 1)^28^ (Suppl. Fig. 3C). Hierarchical clustering of the treatment data revealed three different drug response clusters corresponding to sensitive, intermediate, and resistant TOs (Fig. 3B). Interestingly, despite their clustering in the intermediate or resistant groups, most of these TOs seemed to be sensitive to the HDACi panobinostat and MELKi OTSSP167. Higher-grade MGMs did not seem to be associated with an increased resistance since they were equally distributed among the three groups (Fig. 3C, *P* = 0.72, one-way ANOVA). Different drug responses based on WHO grade were only observed for OTSSP167, with an increased resistance for WHO°3 tumors (Suppl. Fig. 3D). Median IC_50_ values of both drugs panobinostat and OTSSP167 were 10 nM, and thus the lowest concentration tested (Fig. 3D, panobinostat interquartile range: 10-201.9 nM; OTSSP167: 10-48.38 nM). When comparing the median IC_50_ from cell lines and TOs, only panobinostat, OTSSP167, foretinib, and pacritinib showed similar efficacy in both models (ratio 0.4 to 1.4), whereas the other drugs displayed much higher IC_50_ values in TOs (ratio 3.5 to 741), providing evidence for an additional impact of the tumor microenvironment and thus for the potential superiority of the 3D TO model to reflect drug responses of patient tumors. Despite different proposed targets for panobinostat and OTSSP167, AUCs correlated well (Pearson’s correlation r = 0.75, *P* < 0.001; Suppl. Fig. 3E), suggesting a similar vulnerability. Interestingly, *NF2*-wildtype tumors were more sensitive to panobinostat treatment (*P* = 0.039, Student’s t-test; Fig. 3E). Treatment with either OTSSP167 or panobinostat also induced apoptosis in tumor organoids from high-grade MGMs (Fig. 3F&G).

**Fig. 3:**
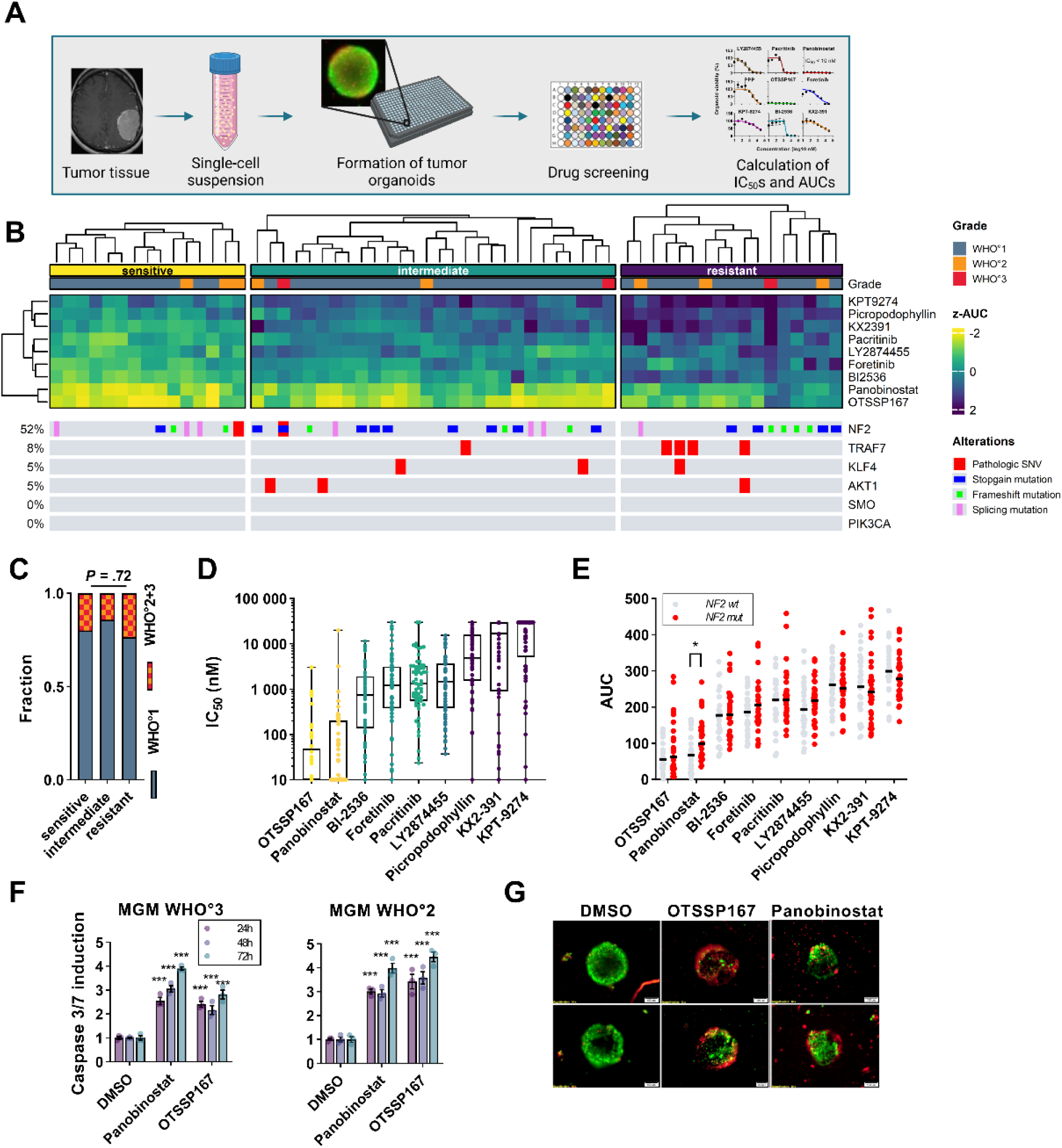
Drug screening in tumor organoids reveals sensitivities to HDACi panobinostat and MELKi OTSSP167. **(A)** Workflow of tumor organoid based drug screening. **(B)** Heatmap of drug responses (z-AUC) from 60 patients. K-means clustering revealed sensitive, intermediate and resistant subgroups. For mutational analysis, only commonly recurring mutations in meningioma were included. This included *AKT1^E17K^*, *KLF4^K409Q^*, *SMO^L412F^* or *SMO^W535L^* alterations and for *PIK3CA* any mutation in codons 542, 545 and 1047. **(C)** Sensitive, intermediate or resistant clusters are independent of WHO grade (Chi-square test). **(D)** Depicted are IC_50_ values for each drug in all 60 cases. OTSSP167 and panobinostat present with the lowest median of 10 nM. **(E)** Comparison of drug responses (AUCs) between *NF2*-wildtype and mutated tumors by Student’s t-tests. **(F)** Tumor organoids from a grade 2 and grade 3 meningioma were treated with panobinostat or OTSSP167 at 1 µM for 24, 48, 72 hrs. Caspase-Glo 3/7 assay was used to determine caspase 3/7 induction. Data was normalized to the control. Student’s t-tests were performed to test significance. **(G)** TOs were treated with 1 µM of panobinostat or OTSSP167 for 24 hrs. Live (green)/dead (red) staining illustrates TO killing.

In summary, we were able to detect robust and reliable drug responses in the majority of meningioma cases for panobinostat and OTSSP167 using a large series of patient-derived TOs.

### Panobinostat’s efficacy is mediated through dual inhibition of HDAC1/2

To validate the known targets of the two most sensitive inhibitors OTSSP167 (MELKi) and panobinostat (HDACi), we treated TOs with additional inhibitors for the same target, and in addition performed RNA interference (RNAi) experiments in MGM cell lines and TOs.

First, we used two additional MELK inhibitors JNJ-47117096 and MELK-8a, of which the latter has been reported to have higher specificity for MELK than OTSSP167^29^. However, treatment of TOs from four patients with JNJ-47117096 and MELK-8a showed no evidence of antineoplastic effects within the same concentration range (Fig. 4A). No reduction of viability below 50% was observed for both drugs up to 10 µM. In line with this finding, neither knockdown of *MELK* nor its downstream effector *EZH2* had an impact on the proliferation of four MGM cell lines (Fig. 4B&C, Suppl. Fig. 4A&B), indicating that OTSSP167 does not mediate its antineoplastic effects through the MELK/EZH2-axis in MGMs.

**Fig. 4:**
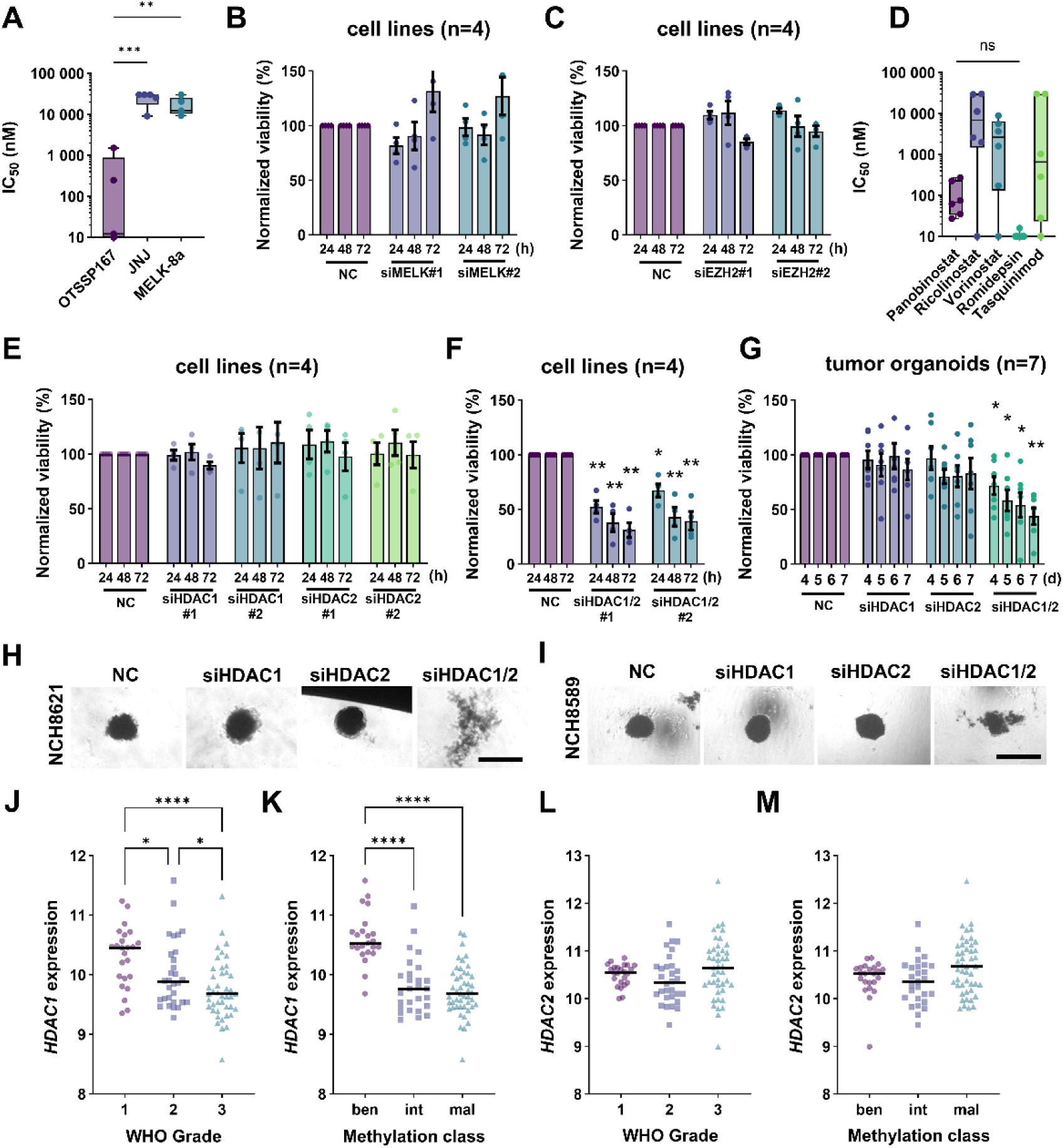
Panobinostat’s efficacy is mediated through dual HDAC1/2 inhibition *in vitro* and *ex vivo*. **(A)** TOs generated from five meningioma patients were treated with the three MELKi JNJ, MELK-8a, and OTSSP167 at half-logarithmic concentrations ranging from 10 nM to 30 µM for 72 h. Dose-response curves were established and corresponding IC_50_ values were calculated. JNJ and MELK-8a failed to demonstrate any effect in tumor organoids, indicating OTSSP167 conveys its effect other than through MELK inhibition. **(B)** RNAi-mediated knockdown experiments for *MELK* or **(C)** its downstream target *EZH2* failed to demonstrate any impact on viability in four meningioma cell lines. Data are presented in mean±SEM. **(D)** TOs from six patients were treated with five different HDACis. Only selective HDAC1/2i romidepsin elicited similar effectiveness compared to panobinostat. **(E)** RNAi-mediated knockdown experiments in four meningioma cell lines fail to demonstrate single knockdown efficacy of *HDAC1* or *HDAC2*. **(F)** Double RNAi-mediated knockdown decreases viability. **(G)** Data was verified in tumor organoids from seven patients. Student’s t-tests were performed comparing double and single knockdowns. **(H&I)** Representative light microscope pictures of two cases transfected with siRNA on day 7. Double knockout altered the morphology. Bar represents 1 mm. **(J)** *HDAC1* mRNA levels from meningioma tissue samples (*n*=97) grouped by WHO grade. **(K)** mRNA expression levels of *HDAC1* grouped by methylation classification. Statistical significance was tested with one-way ANOVA followed by a *post hoc* Dunnett’s multiple comparison test. **(L)** *HDAC2* expression based on WHO grade or **(M)** methylation class. *, *P* < 0.05; **, *P* < 0.01; ***, *P* < 0.001; ****, *P* < 0.0001. NC: negative control.

The HDAC family consists of 18 genes grouped into four classes (I-IV) and two subclasses (IIA/B)^30^. To further determine whether the pan-HDACi panobinostat exerts its effects through a distinct class, we selected inhibitors covering most HDAC classes (ricolinostat, vorinostat, romidepsin, and tasquinimod) and treated TOs from six patients. From those, only romidepsin demonstrated similar efficacy compared to panobinostat (romidepsin vs. panobinostat, *P* = 0.98, Fig. 4D). Romidepsin is considered as a highly selective HDAC1/2 inhibitor^31^. Unexpectedly, single-gene knockdown of *HDAC1* or *HDAC2* did not affect the proliferation of MGM cell lines (Fig. 4E; Suppl. Fig. 4C&D). However, HDAC1/2 are homologs, and there is evidence that the loss of HDAC1 is compensated by HDAC2 and vice versa^32,33^. In line with this assumption, *HDAC1/2* double knockdown decreased viability to 35% in all four cell lines tested (Fig. 4F; Suppl. Fig. 4E). To further substantiate these findings, the impact of RNAi-mediated depletion of HDAC1/2 was validated in MGM TOs from seven patients. In line with the previous findings, double knockdown of *HDAC1/2* in TOs reduced the viability to 43% on day 7 (Fig. 4G, *P* < 0.01) compared to an ineffective single knockdown, indicating vulnerability to loss of HDAC1/2 in meningioma. Microscopical images confirmed a disrupted morphology of HDAC1/2-depleted tumor organoids (Fig. 4H&I).

Next, we interrogated the mRNA expression levels of *HDAC1/2* in transcriptomes of newly diagnosed meningioma tissue samples^25^. Although *HDAC1* mRNA levels were substantially expressed in all tumor samples, the expression levels decreased with WHO grade significantly (Fig. 4J). Similar results were obtained, when the samples were grouped according to their methylation class (MC)^34^. The intermediate and malignant MC group demonstrated significantly lower expression levels than the benign methylation class (Fig. 4K). *HDAC2* expression, however, showed no significant changes within each grade or methylation class (Fig. 4L-M), supporting the compensatory function of *HDAC2*^32,33^.

In summary, panobinostat’s efficacy in MGMs seems to be attributed to the combined inhibition of HDAC1/2.

### Panobinostat treatment reduces tumor growth *in vivo*

To confirm the antimeningioma effect of panobinostat and OTSSP167 *in vivo*, we generated an intracranial xenograft model by injecting anaplastic meningioma cells (NCH93) in the right frontal lobe of NMRI/nu mice. One week after tumor inoculation, mice were randomized and treated intraperitoneally with either panobinostat, OTSSP167 or vehicle control for 5 consecutive days for three weeks followed by discontinuation of the therapy (Fig. 5A). Treatment with panobinostat or OTSSP167 extended median survival from 34 to 42 days (*P* = 0.033, log-rank) and to 44 days (*P* = 0.004, log-rank), respectively. No difference was observed between the two treatment groups (*P* = 0.28, log-rank). In line with the survival data, the proliferation marker Ki-67 was markedly reduced in tumors of treated mice (*P* < 0.001, Fig. 5B&C). Overall, these data suggest a further clinical exploration of panobinostat and OTSSP167 for the treatment of clinically aggressive meningiomas. Because only panobinostat had been FDA- and EMA-approved, we administered panobinostat to a patient suffering from therapy-refractory anaplastic meningioma. DNA methylation profiling and panel sequencing from the tumor demonstrated complex chromosomal aberrations including losses of chromosomes 1p, 4, 6q, 11, 14q, and 17p, and *TP53*, 22q, and homozygous *CDKN2A/B* deletion (Fig. 5D). Panel sequencing revealed nonsynonymous mutations in polybromo-1 (*PBRM1),* chromodomain helicase DNA binding protein 7 (*CHD7),* neurogenic locus notch homolog protein 2 (*NOTCH2),* and DNA polymerase epsilon 2, accessory subunit *(POLE2)* (Fig. 5D). Of note, HDAC1/2 are binding partners of CHD7 and in close relationship to PBRM1, both play an important role in the chromatin remodeling complex^35,36^. Tumor organoids were generated from the tumor of the same patient and treated with the nine best-performing drugs (Fig. 5E). Interestingly, panobinostat and OTSSP167 treatment resulted in flat dose-response curves with 0% viability at the lowest concentration tested (10 nM), which is below panobinostat’s peak serum concentration of 82 nM^37^. Supported by our preclinical data, the patient was treated with panobinostat similar to the European Medicines Agency-approved therapy for multiple myeloma patients receiving 20 mg orally three times a week but without a combination with bortezomib and dexamethasone^38^. MRI scans were routinely conducted at three-month intervals. Although the tumor volume increased from 7.91 cm³ to 9.15 cm³ during treatment (+15% or +1.24 cm³), the preceding tumor growth was largely mitigated (+4.26 cm³ within the first three months after surgery; +3.65 cm³ in three months prior to treatment) (Fig. 5F&G). When assuming linear growth, the extrapolated tumor volume would have been 11.57 cm³ in size (linear regression, *R*^2^ = 0.99). The patient tolerated the oral medication well and developed only mild thrombocytopenia. Noteworthy, a tumor-remote secondary lesion increased during treatment, resulting in the discontinuation of the medication. In conclusion, treatment with panobinostat prolonged the survival of tumor-bearing mice and reduced tumor growth rate in a heavily pretreated anaplastic meningioma patient.

**Fig. 5:**
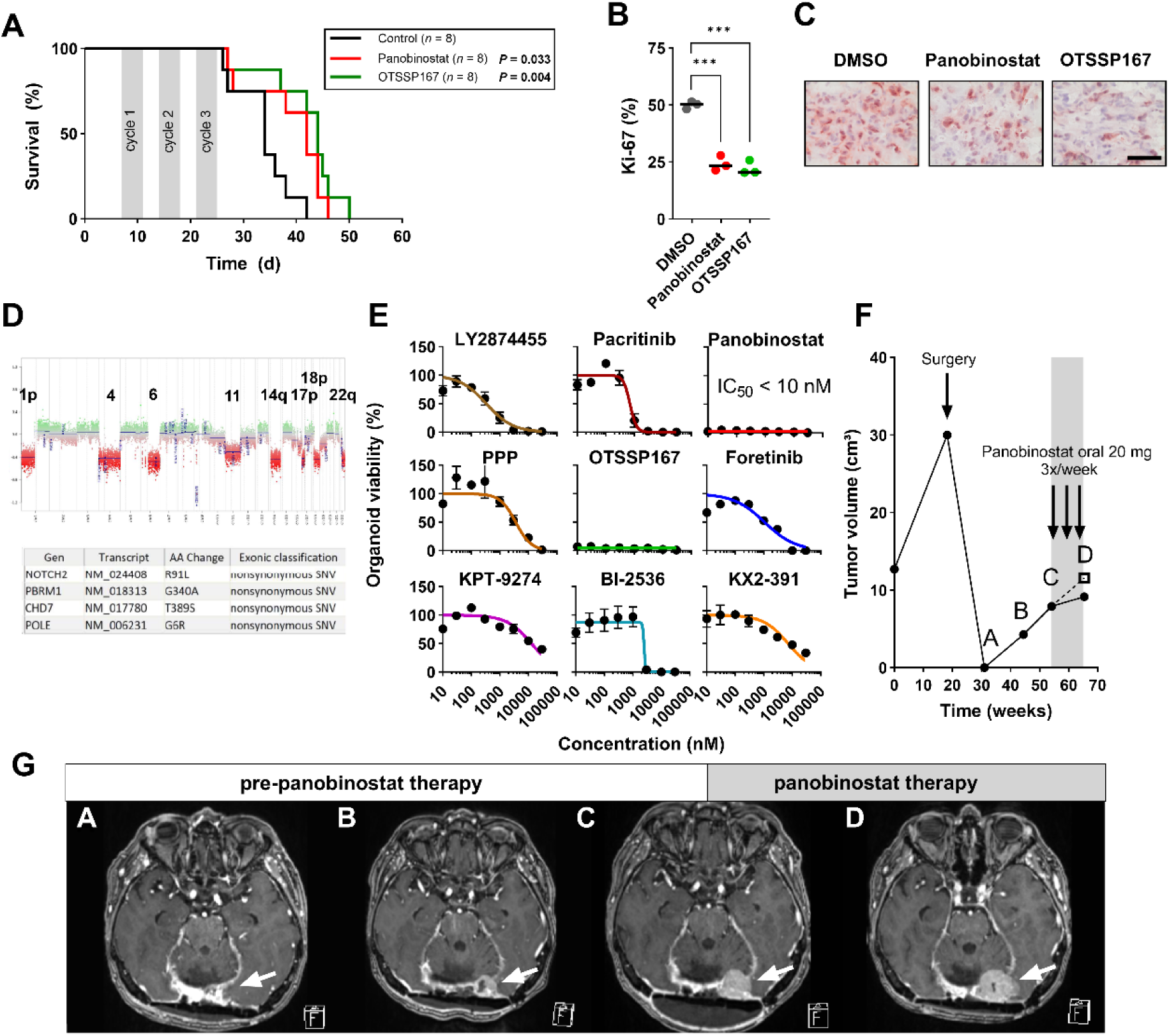
Panobinostat demonstrates efficacy *in vivo*. **(A)** An orthotopic xenograft mouse model was established by injecting NCH93 cells into the right frontal lobe of NMRI/nu mice. After intracranial injection, mice were randomized into treatment groups (*n*=8 per group) and drugs were administered intraperitoneally for three cycles. Treatment was initiated on day 7 with either vehicle (DMSO), panobinostat (20 mg/kg body weight, 5x/week), or OTSSP167 (10 mg/kg body weight, 5x/week). Mice were sacrificed upon signs of neurological symptoms or loss of body weight (>20%). Kaplan-Meier plot shows the survival of mice. Significance was calculated by Log-rank test. **(B)** The Ki-67 expression indicates a significant decrease in the proliferation of treated tumors (*n*=3 per treatment group, one-way ANOVA followed by a *post hoc* Dunnett’s test). **(C)** Immunohistological staining for Ki-67. Bar represents 50 µm. **(D)** Copy number profile and panel sequencing data of a patient suffering from a treatment-refractory malignant meningioma demonstrates complex chromosomal alterations. **(E)** Dose-response curves obtained for 9 selected drugs from the same patient indicate a therapeutic response especially to the HDACi panobinostat. **(F)** When treating this patient with panobinostat, volumetric measurements of contrast-enhancing T1-weighted MRI scans in brain lab software suggest a tumor growth delay as compared to the pre-treatment phase. Dotted line and the square represent the extrapolated tumor volume assuming linear growth from A-C by linear regression model (*R*^2^ = 0.99). Continuous line represents the actual tumor volume. **(G)** Contrast-enhancing T1-weighted MRI images were conducted at three-months intervals (A-D). White arrows point to the tumor. *, *P* < 0.05; **, *P* < 0.01; ***, *P* < 0.001.

### HDACi resistance is conveyed through HDAC8 modulation of the TGFβ-EMT-axis

Because the *in vivo* treatment with panobinostat demonstrated efficacy in improving survival but was not curative, we next explored potential drug resistance. To further elucidate the molecular mechanism of panobinostat resistance, we selected twenty cases with the highest and lowest AUC values of panobinostat and performed bulk RNA-seq from the primary tissues of which TOs were generated (Fig. 6A&B; AUCs: 24 vs 185 or median IC_50_ values 10 nM vs 287 nM, respectively). PROGENγ pathway analysis revealed a significant upregulation of the TGFβ pathway in panobinostat-resistant compared to panobinostat-sensitive tumors (Fig. 6C&D, Estimate = −0.838; *P* = 0.023). TGFβ pathway plays a key role in epithelial-to-mesenchymal transition (EMT), a major player in therapy resistance^39^. In support of our hypothesis, the top result of the hallmark gene set enrichment analysis was the upregulation of EMT in resistant tumors (Fig. 6E, *P* = 0.030). To therapeutically exploit this finding, we hypothesized that the additional treatment with TGFβ inhibitor could overcome HDACi resistance. Three TGFβ receptor I inhibitors (TGFβi; galunisertib, LY3200882, and vactosertib) were tested alone or in combination with panobinostat in meningioma cell lines and panobinostat-resistant tumor organoids. Neither TGFβi-mono nor combination therapy showed any impact on cell viability or synergy concluding that the TGFβ pathway is induced downstream of the receptor (Suppl. Fig. 5A&B). Recent evidence suggests that TGFβ pathway is stimulated by HDAC8^40,41^.

**Fig. 6:**
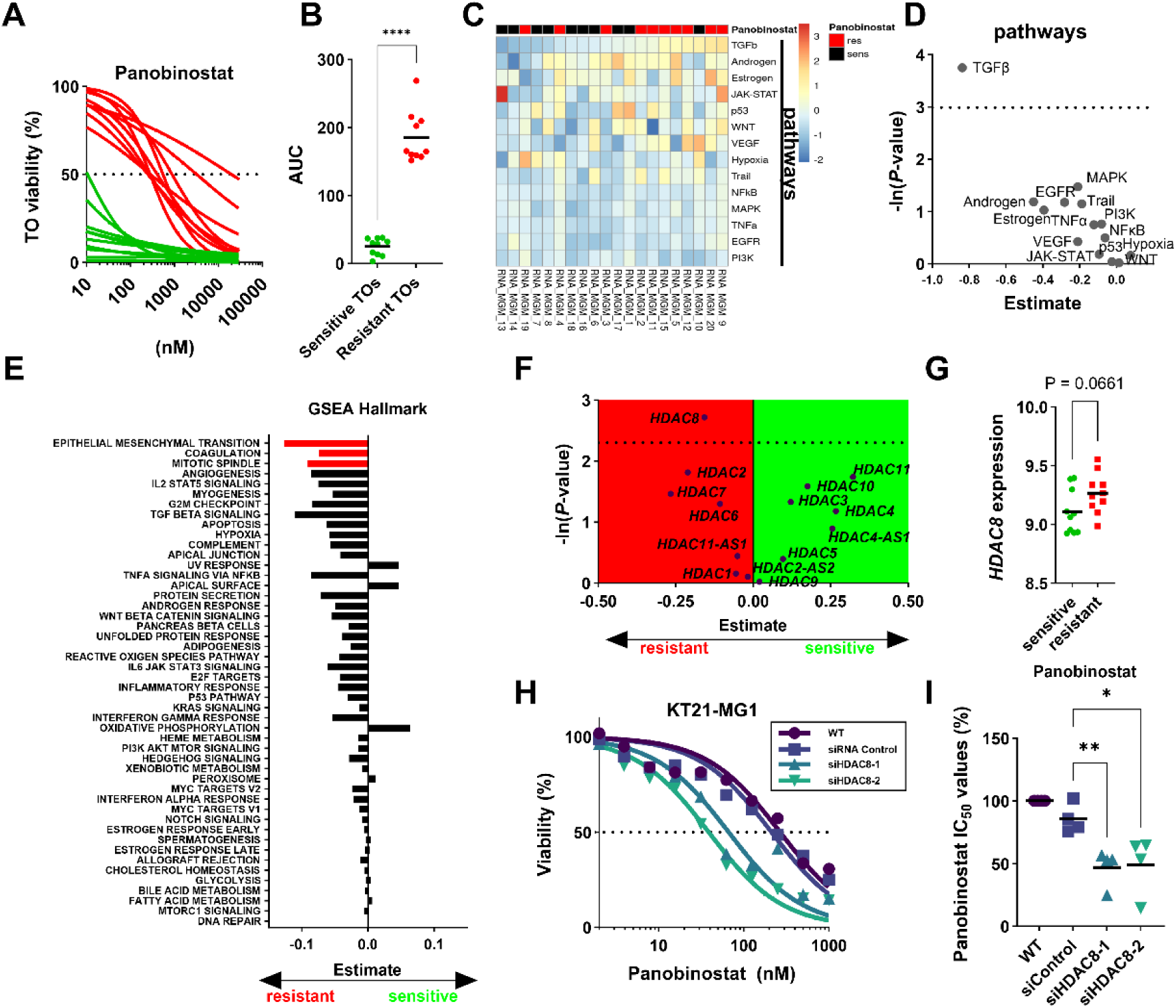
HDAC8 expression mediates intrinsic resistance to panobinostat. **(A)** Dose-response curves of panobinostat depicted from the most sensitive (*n*=10) and resistant (*n*=10) tumor organoids. **(B)** AUCs of panobinostat from sensitive and resistant tumors differ significantly (Student’s t-test). **(C)** Bulk RNA-seq was performed from the corresponding primary tissue samples (*n*=20). PROGENγ inferred pathway activity is presented as heatmap. Res: resistant tumors, sens: sensitive to panobinostat. PROGENγ pathway analysis demonstrates that **(D)** TGFβ pathway is significantly activated in panobinostat-resistant tumors calculated by linear model analysis (estimate: sensitive vs resistant). **(E)** GSEA hallmark analysis demonstrates significant upregulation of EMT, coagulation, and mitotic spindle in resistant tumors. Significant results are marked in red (*P* < 0.05, one-way ANOVA). **(F)** Linear model analysis indicates that *HDAC8* expression might play a role in panobinostat resistance (estimate: sensitive vs resistant). **(G)** *HDAC8* mRNA levels are increased in panobinostat-resistant tumors (Student’s t-test). **(H)** Knockdown of *HDAC8* increased sensitivity to panobinostat. DRCs of panobinostat were established using meningioma cells (*n*=4) transfected with two different siRNAs targeting *HDAC8* or control. DRCs of panobinostat derived from KT21-MG1 meningioma cell line are shown as an example. **(I)** Data from all meningioma cell lines were normalized to the untransfected wild-type control (WT) (one-way ANOVA, Dunnett’s post hoc test). *, *P* < 0.05; **, *P* < 0.01; ***, *P* < 0.001; ****, *P* < 0.0001. TO: Tumor organoid.

Therefore, we investigated the differential gene expression of HDAC genes. Of note, only *HDAC8* mRNA was upregulated in resistant tumors (Fig. 6F&G, *P* = 0.066, Estimate = −0.15). We hypothesized that the intrinsic upregulation of *HDAC8* promotes EMT through the TGFβ-axis causing resistance to panobinostat. Therefore, the impact of HDAC8 depletion was tested on the sensitivity of panobinostat in meningioma cell lines. In line with our hypothesis, RNAi experiments resulted in a left shift of the dose-response curve, indicating increased sensitivity towards the drug (Fig. 6H&I). Taken together, we observed that panobinostat resistance seems to be conveyed through HDAC8-TGFβ-EMT-axis and therefore additional targeting HDAC8 might be an attractive option to increase the efficacy of panobinostat. Furthermore, these findings provide insights into the molecular mechanisms underlying panobinostat resistance and highlight the potential of tumor organoids in investigating clinically relevant drug resistance.

## DISCUSSION

The management of aggressive meningiomas remains challenging due to limited treatment options besides surgical removal and radiotherapy^2^. High recurrence rates, especially in high-grade meningiomas, and a lack of effective systemic treatment are responsible for the unfavorable prognosis^3^. Here, we conducted an *in vitro* drug screening with targeted anticancer drugs and discovered nine sensitive inhibitors. Subsequent validation was performed on a highly standardized tumor organoid model recapitulating meningiomas histologically and genetically. Personalized drug screening on the so far largest cohort of meningioma TOs of 60 patients revealed substantial antitumor effects of the HDACi panobinostat and the MELKi OTSSP167 in the majority of cases. Panobinostat’s effect was mainly mediated through HDAC1/2 inhibition, whereas for OTSSP167 off-target effects are likely. Both drug treatments prolonged the survival of mice. In addition, panobinostat monotherapy delayed the tumor growth of a therapy-refractory anaplastic meningioma patient. In this very advanced case, however, therapy was not curative. Lastly, we used our novel TO model to decipher a potential intrinsic panobinostat resistance by providing substantial evidence that this was mediated by a HDAC8-TGFβ-induced EMT-like mechanism.

In this work, we highlight the power of a multi-step drug screening workflow aimed at identifying potential therapies for aggressive meningiomas. Initially, we started with an extensive screening process using four aggressive meningioma cell lines to filter out less sensitive drugs, resulting in the elimination of 92% of the 107 candidates. Then, the top nine drugs underwent further investigation in freshly prepared meningioma tumor organoids from as many as 60 patients, encompassing the largest cohort of tumor organoids studied to date, and covering most histological and mutational characteristics of meningioma. Only two of the nine drugs revealed substantial sensitivity in TOs, indicating the superiority of the TO model by maintaining the tumor microenvironment and patient individual mutational and transcriptomic landscape in tissue-like 3D structures. Finally, the efficacy of the two most promising drugs, OTSSP167 and panobinostat, was confirmed in an orthotopic xenograft model and complemented for panobinostat through the treatment of one patient suffering from therapy-refractory meningioma. This comprehensive workflow - from cell lines to tumor organoids and mouse models to a patient – suggests the HDACi panobinostat as promising drug for treatment of aggressive meningioma and identified a potential intrinsic panobinostat resistance mechanism.

HDACs and histone acetyltransferases (HATs) orchestrate gene expression through de- or acetylation of histones and non-histone proteins^30^. By modifying histone charges, accessibility to DNA for transcription factors is altered resulting in epigenetic reprogramming of the cell which is now recognized as a hallmark of cancer^42^. In our study, pan-HDACi panobinostat was highly effective in meningioma cell lines and next-generation tumor organoids with a median IC_50_ of 10 nM for the latter independent of WHO grade suggesting a wide range of applications. Besides us, only a few other groups have previously tested HDACi in meningioma cell lines, primary cells, and patient-derived orthotopic xenograft mouse models^17,43–45^. Among them, HDACi panobinostat and romidepsin demonstrated one of the highest antimeningioma effects with IC_50_ values in the lower nanomolar range comparable to our current and previous results^17,43–45^. We further identified that the effects of panobinostat were mainly attributed to simultaneous inhibition of HDAC1/2 since single knockout had no impact on meningioma proliferation which most likely is due to the ability of HDAC1/2 to compensate the lack of each other^32,33^. Still, the question remains why meningiomas are so remarkably sensitive to HDAC1/2 inhibition? Recent data have shown that HDAC1 is recruited within super-enhancers marked by acetylated H3K27 and BRD4 functioning as a transcriptional activator for survival and progression in chronic lymphocytic leukemia^46^. Upon HDAC1 inhibition, expression of related genes is effectively abolished^46^. Similarly, Marié *et al.* suggested that acetylated histones bind BRD4 which in turn recruits the transcription machinery^47^. HDACi leads to the accumulation of acetylated histones and therefore diminishes the pool of free BRD4 and decreases gene expression^47^. Our microarray and methylation data suggest that especially *HDAC1/2* among the HDAC family members are highly expressed. However, expression of *HDAC1* mRNA is highest in grade 1 meningiomas and decreases with malignancy. Similar to the *in vitro* RNAi experiments, the reduction of *HDAC1* mRNA levels might be compensated by *HDAC2*^32,33^.

The pan-HDACi panobinostat is an orally bioavailable drug and received approval in the US and Europe in combination with bortezomib and dexamethasone for patients with relapsed and refractory multiple myeloma. The peak plasma concentration of panobinostat attains 82 nM^37^ indicating that effective concentrations (IC_50_<82 nM) can be achieved in 70% (n=42/60) of all tested meningioma cases in this study. Moreover, a patient with a heavily pretreated recurrent anaplastic meningioma was treated with 20 mg panobinostat orally three times a week. Although tumor still grew during panobinostat therapy, growth was remarkably delayed compared to the pretreatment period. Taking into consideration that the patient tolerated this treatment well, we propose that treatment at a less advanced stage of the disease, an increased dose or shortening the dose intervals could enhance the observed effects in future clinical studies. Furthermore, a search for more effective combinations is warranted to maximize the effects of panobinostat. In summary, panobinostat is a highly effective drug in meningiomas at nanomolar levels, it causes its antimeningioma effects through inhibition of HDAC1/2 and deserves further clinical evaluation.

Drug resistance is an emerging problem in targeted therapy and overcoming resistance is key to prolonging survival of patients^48^. The application of tumor organoids in exploring resistance mechanism is a novel field^49^. The current analysis clearly shows its suitability to decipher drug resistance by including bulk RNA sequencing of primary tumors sensitive and resistant to the drug. Using this approach, we discovered that HDAC8, TGFβ-pathway, and EMT-like mechanism are upregulated in panobinostat-resistant tumor samples. Recent evidence supports that HDAC8 induces the expression of downstream TGFβ-pathway genes, which in turn activates the EMT process^39–41^. In line with this hypothesis, downregulation of HDAC8 in meningioma cell lines increased their sensitivity to panobinostat. In cancer cells, HDAC8 plays a multifunctional role by acting on histones and non-histone substrates including p53, SMC3, and estrogen-related receptor alpha (ERRα)^50^. It stimulates tumor growth, metastasis, enhances cell proliferation, suppresses apoptosis, and activates EMT^50^. Furthermore, HDAC8 is involved in immune evasion and drug resistance^50^. Recently, several studies implicated HDAC8 to withstand chemotherapeutic drugs like paclitaxel^41^, temozolomide^51^, doxorubicin^52^, or daunorubicin^53^, making it an interesting target to overcome panobinostat resistance.

In conclusion, we provide strong *in vitro*, *in vivo*, *ex vivo*, and patient evidence for the efficacy of the HDACi panobinostat to treat clinically aggressive meningiomas and uncovered a potential intrinsic resistance mechanism by activation of the HDAC8-TGFβ-EMT axis.

## MATERIALS AND METHODS

### Cell lines

The hTERT-transformed meningioma cell line Ben-Men-1^54^ (DSMZ, Braunschweig, Germany) and the anaplastic meningioma cell lines NCH93^55^, IOMM-Lee^56^ (ATCC), and KT21-MG1^57^ (kindly provided by Prof. Mawrin, University of Magdeburg) were cultured in Dulbecco’s minimal Eagle’s medium (DMEM, Gibco) supplemented with 10% fetal calf serum (Sigma-Aldrich), 2% L-GlutaMAX (Sigma-Aldrich), and 1% Penicillin/Streptomycin (Sigma-Aldrich) at 37°C in a humidified environment with 5% CO_2_ atmosphere. Mycoplasma contamination was excluded by 4’,6-diamidino-2-phenylindole staining (Roche, Basel, Switzerland). Cell lines were authenticated by STR DNA profiling analysis (Leibniz Institute DMSZ, Braunschweig, Germany).

### Targeted anticancer drug library

An *in silico* analysis was performed based on the microarray data of 28 cases of WHO grade 3 meningioma previously published by our group^25^. For more than 20,000 genes expressed in malignant meningiomas, we looked for the availability of specific inhibitors using Selleck Chemical’s website (https://www.selleckchem.com/). 442 genes fulfilled this requirement. Subsequently, the NIH Clinical Trial website (https://www.clinicaltrials.gov/) was used to determine the clinical trial status of all inhibitors. Only inhibitors were included which were currently used in clinical trials or were FDA-approved. In total, 182 inhibitors targeting 72 genes were identified. Furthermore, we excluded FDA-approved drugs that overlapped with the AODVI drug library which was previously studied in our lab^17^, resulting in 143 inhibitors that targeted 46 genes. Lastly, a maximum of three inhibitors per gene were selected for further investigation, which led to a drug library containing 107 inhibitors, targeting 46 genes (Supplementary Table 1).

### Large-scale drug screening *in vitro*

NCH93 and IOMM-Lee cells were seeded at a density of 4,000 cells/well in 96-well plates. On the next day, cells were treated with each compound (*n*=107) at a single dose (2.5 µM) in triplicates for 48 h. The viability was assessed by CellTiterGlo 2.0 (Promega, Madison, WI, USA) according to the manufacturer’s recommendations. The viability was normalized to DMSO control-treated cells. For the second step of drug screening, NCH93, IOMM-Lee, Ben-Men-1, and KT21-MG1 were seeded at a density of 4,000 cells/well in 96-well plates. On the next day, cells were treated with each compound (*n*=33) at six concentrations ranging from 0.1 to 10,000 nM for 48 h. Cell viability was assessed by CellTiter-Glo 2.0 (Promega). The IC_50_, AUC, and E_max_ values were calculated using nonlinear regression in GraphPad Prism. AUC and E_max_ data were z-transformed.

### Crystal violet assay

Meningioma cells were treated with drugs (*n*=9) at 24 doses ranging from 0.000275 to 10,000 nM for 48 h. After the removal of medium, cells were washed with 100 μL DPBS (Gibco, New Hampshire, USA) once. Subsequently, 50 μL of 0.5% crystal violet (Sigma-Aldrich) was added into wells, and cells were incubated for 15 min at room temperature (RT) on a shaker. Next, crystal violet was removed and cells were washed with aqua bidest (B.Braun). Plates were left to dry overnight in a fume hood. On the following day, 200 μL of methanol (Sigma-Aldrich) was added to dissolve crystal violet. The optical density was recorded at 555 nm using a microplate reader (Infinite F200 pro, Tecan GmbH, Groedig, Austria). IC_50_ values were calculated as described above.

### BrdU proliferation assay

Cell proliferation was quantified using the Cell Proliferation ELISA, BrdU Kit (Roche, Basel, Switzerland). NCH93, Ben-Men-1, IOMM-Lee, and KT21-MG1 cells were cultured 4,000 cells were seeded in 96-well plates in 100 µL medium. On the next day, cells were treated with each compound (n=9) at 5xIC_50_ derived from the crystal violet assay. After 24, 48, and 72 h, 10 µL BrdU labeling solution was added to the cells (final concentration of 10 µM BrdU) and incubated for two hours. Next, the medium was removed from the cells by tapping off. Cells were fixed with 200 µL per well FixDenat and incubated for an additional 30 min at RT. FixDenat was removed by tapping off and 100 µL anti-BrdU-POD working solution per well was added and incubated for 90 min at RT. The antibody conjugate was removed and wells were rinsed three times with 200 µL PBS. Thereafter, 100 µL substrate solution was added and incubated for an additional 10 min at RT. Luminescence was measured using Microplate Reader Tecan Infinite PRO. Data were normalized to the DMSO control group.

### Cell cycle analysis

Meningioma cells were seeded at a density of 3×10^5^ in 6-well plates. Thereafter, cells were treated with each compound at 5xIC_50_ or DMSO control for 24 or 48 h. Next, cells were harvested and fixed with 85% ice-cold ethanol (Roth). After the incubation period and a washing step, 5 μL RNAse A (Lucigen) was added to the cells and incubated for 30 min. Before measurement, propidium iodide (PI) (Sigma-Aldrich) was added at a final concentration of 50 µg/mL. DNA content was analyzed by LSR II flow cytometer (BD Biosciences, San Jose, CA, USA) and cell cycle distribution was analyzed using the software FlowJo (FlowJo, Oregon, USA).

### Caspase 3/7 assay

The Caspase-Glo 3/7 Assay (Promega) was used to measure apoptosis in meningioma cells. Cells were seeded at a density of 4,000 in 96-well plates. On the next day, cells were treated with each compound (n=9) at 5xIC_50_. After 24, 48, and 72 h, equal volumes of Caspase-Glo 3/7 Assay reagent were added to each well according to the manufacturer’s recommendation. Luminescence was measured using Microplate Reader Tecan Infinite PRO. Data were normalized to DMSO control.

### Tumor samples and molecular analyses

The study was approved by the institutional review board. Informed consent was obtained from all patients. Fresh tumor material was divided for preparing single cell suspensions and for cryoconservation until further processing. Meningioma tissue samples and therefrom derived tumor organoids were profiled by targeted panel sequencing to identify meningioma-typical aberrations. The panel (NPHDA2022) contained genes reported to be frequently mutated in brain tumors including MGMs as described previously^58^. Sequencing was done by applying a custom hybrid capture approach (Agilent Technologies, CA, USA) on a NextSeq500 instrument (Illumina). For single nucleotide variant and InDel calling, SAMtools mpileup and Platypus were utilized. The panel was designed to assess the frequency of known brain tumor mutations rather than novel mutational events. Meningioma methylation classes were determined by random-forest classifier trained for established meningioma methylation classes benign, intermediate and malignant as described previously^34^. For *HDAC1/2* expression analyses, we used our previously published microarray data set^25^. RNA input libraries were constructed according to manufacturer’s instructions (TruSeq Stranded Total RNA, Illumina, San Diego, CA), and sequenced on Illumina NovaSeq 6000, 100bp PE reads in the DKFZ genomics and proteomics core facility. Reads were aligned against GRCh38.p7, using STAR v. 2.6.0c^59^. Count data was vst normalized with the DESeq2 package^60^. Tumor pathway activity was inferred using PROGENy^61^. Heatmaps were generated using the ComplexHeatmap and pheatmap packages in R.

### Establishment of patient-derived meningioma tumor organoids

Fresh meningioma tissue was mechanically minced into small pieces and digested by the Tumor Dissociated kit (TDK, Milteny) in the GentleMACS dissociator. The generated single-cell suspension was filtered through cell strainers and incubated in 5 ml Red Blood cell lysis buffer (150 mM NH_4_Cl, 10 mM KHCO_3_, 0.1 mM EDTA) for 5 min at RT. Next, 20 ml PBS was added and the suspension was centrifuged at 300g for 10 min. The cell pellet was resuspended in a 5 ml cell culture medium. Thereafter, cells were incubated with NanoShuttle (1 µL per 10^4^ cells, Greiner Bio-One) overnight at 37°C. On the next day, 1,250 to 100,000 cells per well were seeded in anti-adhesive 96- or 384-well plates and put on magnets for 30 min to allow the formation of tumor organoids. Images of tumor organoids were taken for 14 consecutive days. The diameter of TOs was measured using ImageJ (1.8.0). The viability of TOs was daily measured using CellTiter-Glo 3D (Promega) and normalized to day 0.

### Large-scale tumor organoid drug screening

Tumor organoids from tumors of 60 meningioma patients were generated in anti-adhesive 384-well plates. To ensure the sufficient representation of low abundance cells 25,000 cells/TO were chosen. On day 3, TOs were treated with the nine compounds at eight semi-logarithmic doses ranging from 10 to 30,000 nM. After 72 h, equal volumes of CellTiter-Glo 3D were added to the wells. The plates were then placed on a shaker for 5 min. After incubation for 25 min at RT, a plate reader (Infinite F200 Pro, Tecan GmbH, Groedig, Austria) was used to record luminescence. Data were normalized to DMSO controls and IC_50_ and AUC values were calculated as described above.

### Live/Dead staining

Treated or non-treated tumor organoids were incubated with an equal volume of the live/dead cell imaging kit (Invitrogen). To each well and incubated for 15 min at RT. Images were taken with the fluorescence microscope (Zeiss, Oberkochen, Germany) and cellSens Dimension software was used for analysis.

### DNA/RNA isolation

Only tissue samples with a vital tumor cell content >60% as determined on H&E-stained slides from each tissue by a board-certified neuropathologist were eligible for RNA/DNA extraction (Department of Neuropathology, University Hospital Heidelberg, Germany). Total RNA was extracted from tissues using the AllPrep Kit (Qiagen) according to the manufacturer’s instructions. RNA and DNA from tumor organoids were extracted with RNeasy Mini Kit (Qiagen) or DNeasy Blood & Tissue Kit (Qiagen). RNA integrity was assessed by the Agilent 2100 Bioanalyzer and quantified by NanoDrop ND-1000 spectrophotometer (Thermo-Scientific, Waltham, MA, USA) and then stored at −80°C until further analysis.

### Immunohistochemical staining

Acetone-fixed cryosections of xenografted mouse tumors were stained with rabbit polyclonal anti-Ki-67 (1:50 dilution, ab15580, Abcam, Cambridge, UK) antibody diluted with DAKO diluent (Agilent) for 60 minutes at RT, and washed three times with PBS-Tween-20 (0.05%). Next, the corresponding secondary antibody (Vector, Burlingame, USA) diluted in goat serum (Vector, Burlingame, USA) and DPBS (Gibco) was added to the slides and incubated for 30 min. After repeated washing steps, an avidin-biotin-complex (Vector, Burlingame, USA) was added to the slides for 30 min. Thereafter, the AEC substrate (Vector, Burlingame, USA) was added to the slides and incubated for 3 min. Finally, slides were counterstained with hematoxylin (Sigma-Aldrich). Ki-67 positive cells were counted in ten high-power fields per slide. Rabbit IgGs served as a negative control.

### Immunofluorescence staining

Staining was performed on acetone-fixed cryosections (5-8 µm) of meningioma tumor organoids. Anti-EMA antibody (1:100 dilution, mouse monoclonal IgG, BioLegend), anti-fibronectin antibody (1:200 dilution, rabbit monoclonal IgG, Agilent), and Anti-CD68 antibody (1:25 dilution, mouse monoclonal IgG, Agilent) were diluted in DAKO diluent (Agilent). Slices were incubated with primary antibodies at RT for 60 min and washed three times with PBS-Tween-20 (0.05%). Thereafter, the slides were incubated for 60 min with the corresponding secondary antibody (1:200 dilution, Invitrogen) diluted in DAPI (1:1000 dilution, Invitrogen) and DPBS (Gibco), followed by three washing steps with PBS-Tween-20. Finally, coverslips were mounted with elvanol (Sigma-Aldrich). A fluorescence microscope (Zeiss, Oberkochen, Germany) and cellSens Dimension software were used for imaging and analysis.

### RNAi-mediated knockdown

Meningioma cells were transfected with siRNA against *HDAC1* (Cat. #4427037-s73, 4427038-s74, Invitrogen), *HDAC2* (Cat. #4427037-s6493, 4427038-s6495, Invitrogen), *MELK* (Cat. #4427037-s386, 4427038-s387, Invitrogen), *EZH2* (Cat. #4427037-s4916, 4427037-s4918, Invitrogen) or siRNA negative control (Cat. #4390843, Invitrogen). Lipofectamine RNAiMAX reagent (Cat. #13778030, Invitrogen) was used to deliver siRNAs into the cells. Cells were transfected with a final concentration of 100 nM siRNA and 7.5 µL Lipofectamine RNAiMAX in 1 ml OptiMEM (Gibco) overnight. On the next day, the medium was changed to DMEM supplemented with 10% FCS, 2% GlutaMAX (Gibco), and 1% Pen/Strep, and cells were further used for experiments. Validation of the knockdown was assessed by qRT-PCR. A decrease in mRNA levels of >70% compared to siRNA negative control was used as a threshold for a successful siRNA-mediated knockdown of the target gene.

### Quantitative Real-Time PCR

Equal amounts of total RNA (1 µg) were reverse-transcribed using the Transcriptor First Strand cDNA Synthesis Kit (Roche) with random hexamer primers for one hour at 50°C. qPCR was performed in triplicates on a LightCycler 480 (Roche) using the LightCycler 480 Probes Master and probes from the Universal Probe Library (Roche) as described [www.roche-applied-science.com]. Relative fold changes between the expression of target genes were calculated using the 2^-ΔΔCq method. *GAPDH*, *ACTB*, and *HPRT1* were used as reference genes. Relative expression levels of *HDAC1*, *HDAC2*, *MELK*, and *EZH2* mRNA levels were normalized to the RNAi control samples. The primers used are shown in Suppl. Table S2.

### Orthotopic xenograft mouse model

All experiments were done following the regulations of animal protection and approved by Regierungspraesidium Karlsruhe (#35-9185.81/G-99/21). Five to 6-week-old female NMRI/nu mice (Janvier Laboratory, Le Genest-Saint-Isle, France) received intracranial injections of 1×10^5^ NCH93 cells in 5 µL PBS using a stereotactic device into the right frontal. One week after tumor inoculation, mice were randomly separated into three groups (n=8). Mice were intraperitoneally treated with panobinostat (20 mg/kg, five consecutive days a week), OTSSP167 (10 mg/kg, five consecutive days a week), or vehicle control (DMSO) for three cycles. Animal behavior and weight of the mice were monitored daily. The mice were sacrificed upon showing signs of tumor formation (rough coat, hunching, neurological deficit, and weight loss). The survival time of each mouse was recorded. The tumor-containing mouse were explanted, snap-frozen, and stored at −80°C until further analysis.

### Statistical Analysis

All *in vitro* experiments were performed at least in triplicates, and results were expressed as mean ± SEM. *P*-values were calculated using a two-tailed Student’s t-test, one-way ANOVA, or log-rank test in GraphPad (Ver. 9.5.0). *P*-values < 0.05 were considered significant (* *P* < .05; ** *P* < .01; *** *P* < .001; **** *P* < .0001).

## Acknowledgement

We thank Ronja Trunk, Lisa Petermann, Farzaneh Kashfi, Ilka Hearn, and Melanie Greibich for their excellent technical assistance.

## Author Contributions

conceptualization, G.J., and C.H.-M.; methodology, G.J., J.C., R.W., M.M., F.S., M.K., C.H.-M.; software, G.J., J.C., T.Y., M.K., R.W.; validation, G.J., J.C.; formal analysis, G.J., J.C., M.K., F.S., R.W.; investigation, J.C., G.J., T.Y, C.L., V.B., L.J., P.DT., A.Y., M.S., F.S., M.M., M.K.; resources, C.H.-M., A.U., S.K., A.A., M.B., A.v.D., F.S.; writing, G.J., C.H.M.; visualization, G.J., J.C.; supervision, C.H.-M.; project administration, G.J., C.H.-M..; funding acquisition, C.H.-M., G.J., R.W.

## Data and materials availability

The data that support the findings of this study are available from the corresponding author upon reasonable request.

## SUPPLEMENTARY FIGURES

**Suppl. Fig. 1:**
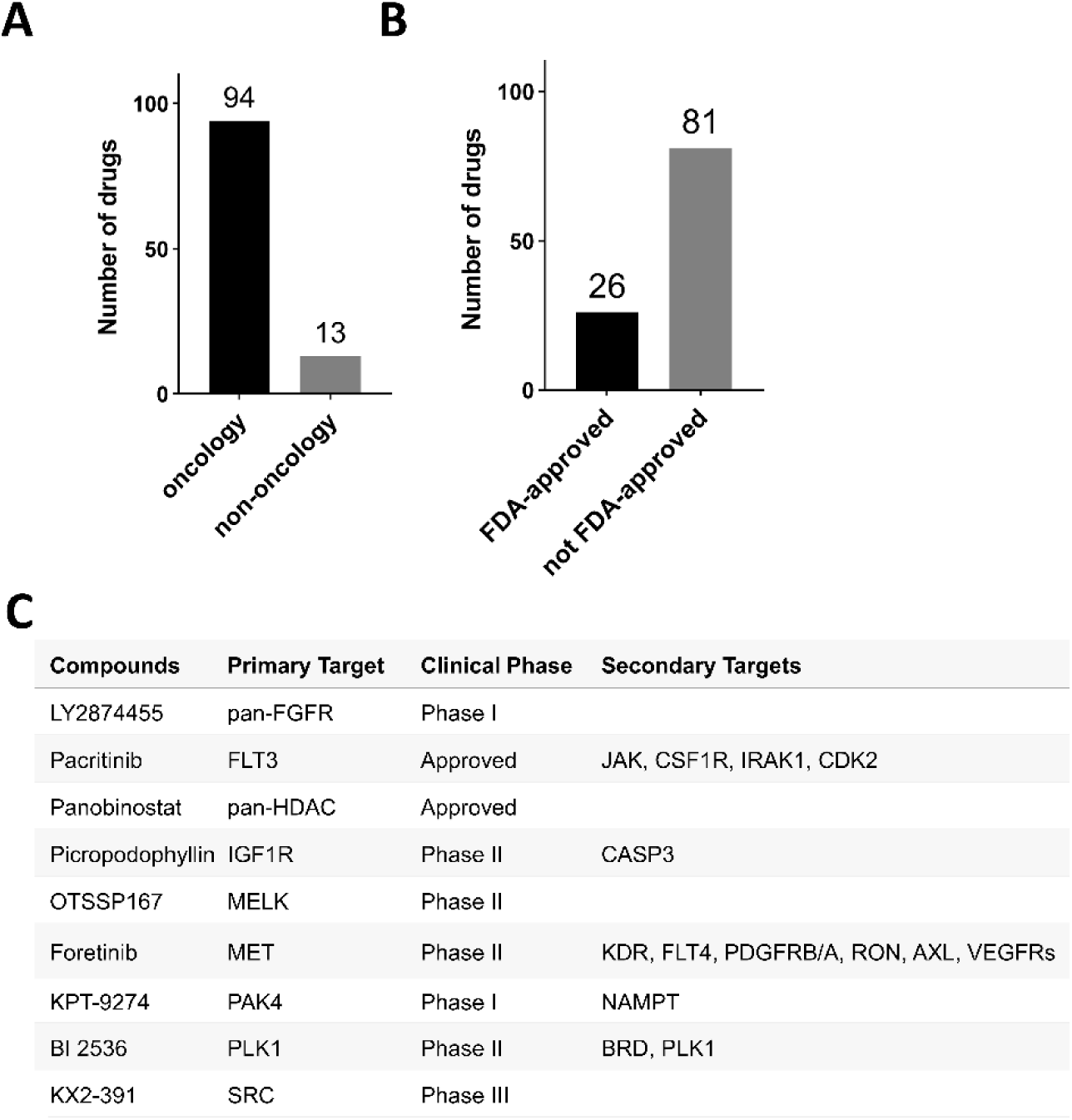
Drug characterization. **(A)** Number of oncology and non-oncology drugs. **(B)** Number of FDA-approved and non-FDA-approved drugs. **(C)** Overview of the top nine drugs, their primary and secondary targets and current clinical development status.

**Suppl. Fig. 2:**
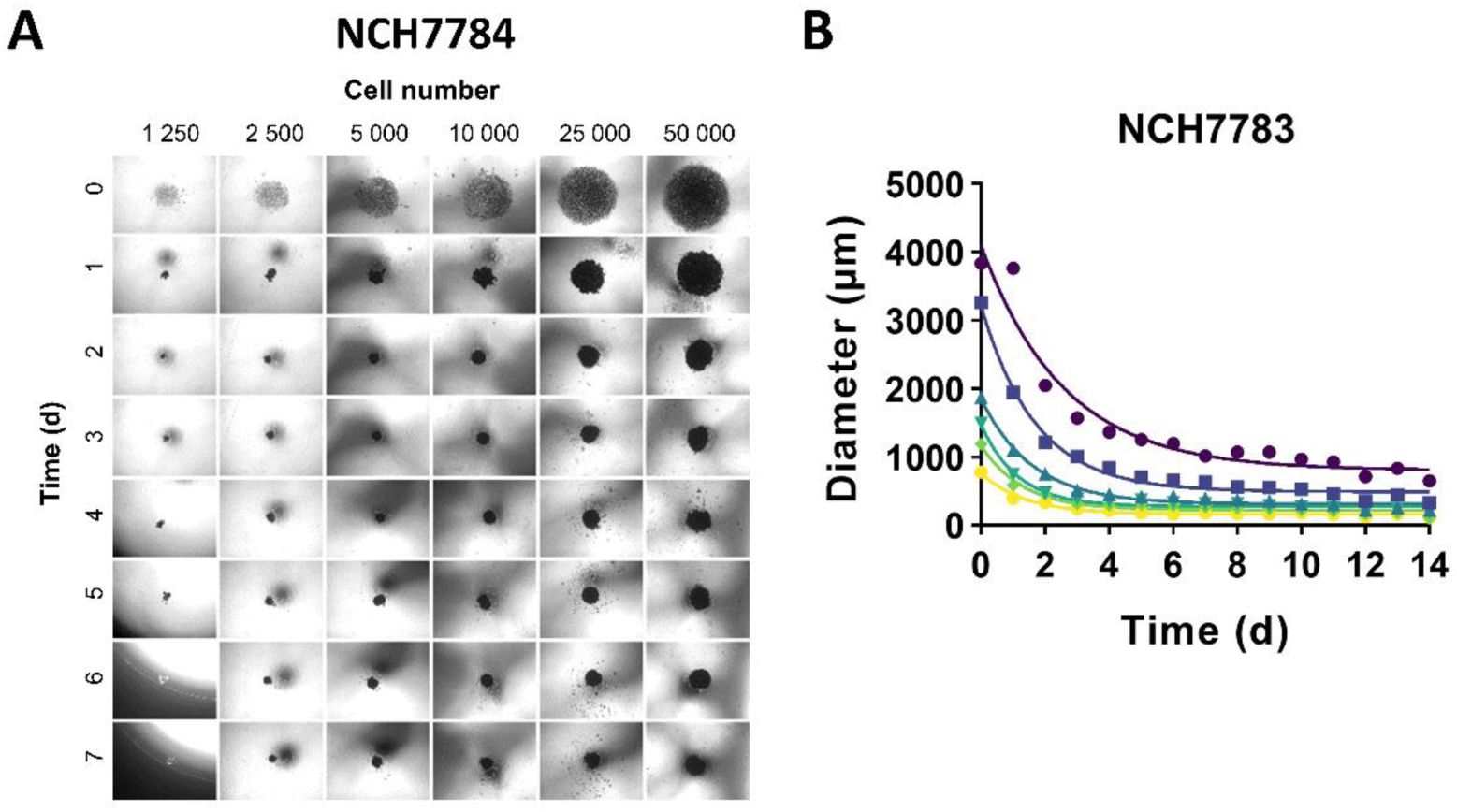
Tumor organoid characterization. **(A)** Depicted are corresponding representative light microscope images of tumor organoids from Fig. 2C. **(B)** The diameters of tumor organoids were measured from another meningioma (NCH7783). Similarly, cells condensate to compact tumor organoids within 2-4 days and stay stable in size thereafter.

**Suppl. Fig. 3:**
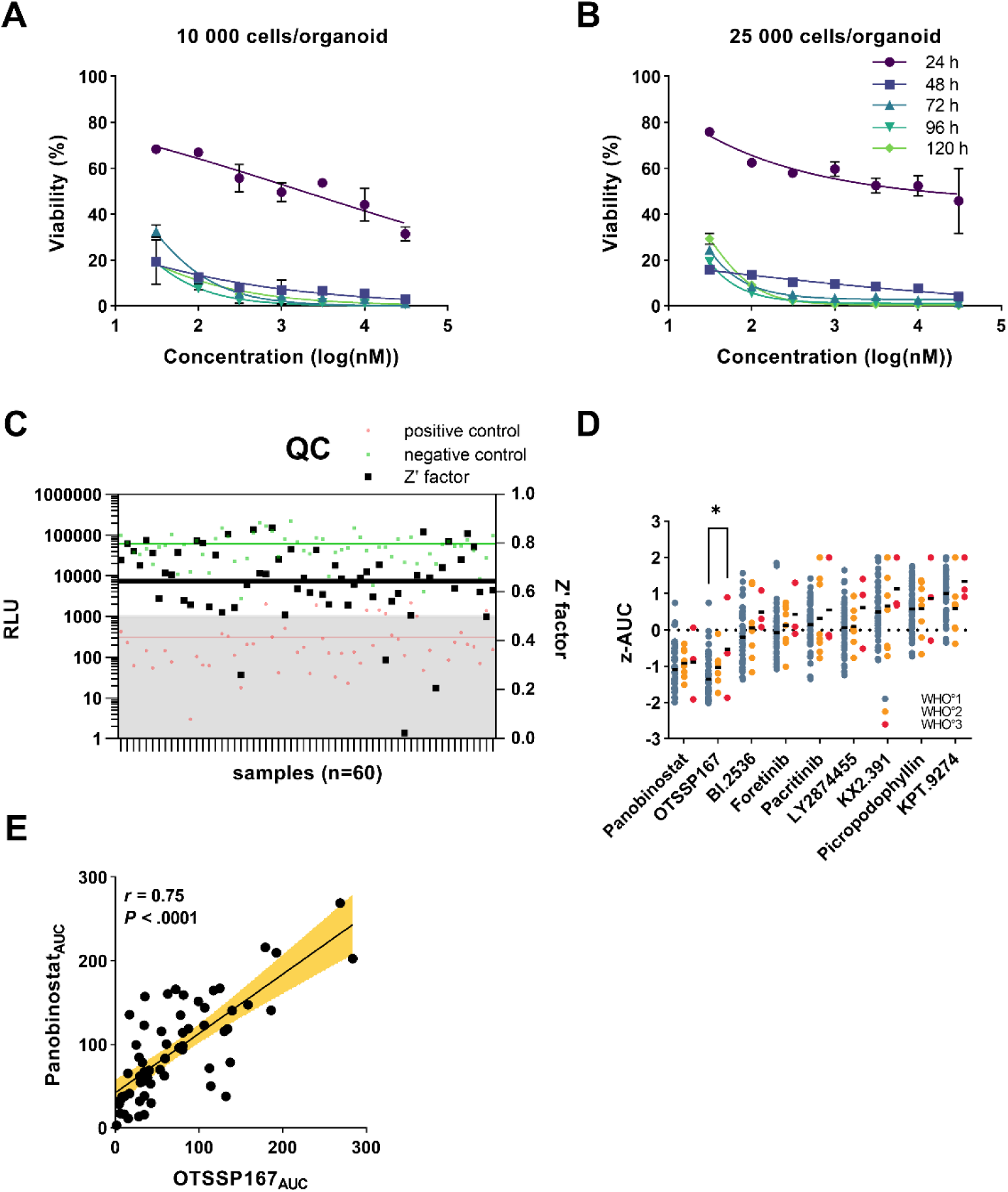
Tumor organoid treatment. **(A)** TOs were generated with 10,000 or 25,000 **(B)** cells/organoid and treated with Panobinostat ranging from 30 nM to 30 µM for 24, 48, 72, 96, and 120 hrs. The viability was assessed by CellTiterGlo3D. Linear regression analysis was performed in GraphPad. Maximum treatment efficacy was achieved after 48 to 72 hrs independent on the organoid size. **(C)** Z’ factor as an indicator for quality control for drug screenings. Z’ factor was calculated from the relative light units (RLU) from negative (DMSO, green) and positive (staurosporin, red) control in every case. The overall Z’ factor was calculated from the average of all individual Z’ factors. (**D)** Comparison of drug responses (z-AUCs) between the WHO grades. One-way ANOVA followed by post hoc analysis was performed to test significance. **(E)** Pearson’s correlation of drug responses (z-AUCs) of HDACi panobinostat and MELKi OTSSP167. *, *P* < 0.05.

**Suppl. Fig. 4:**
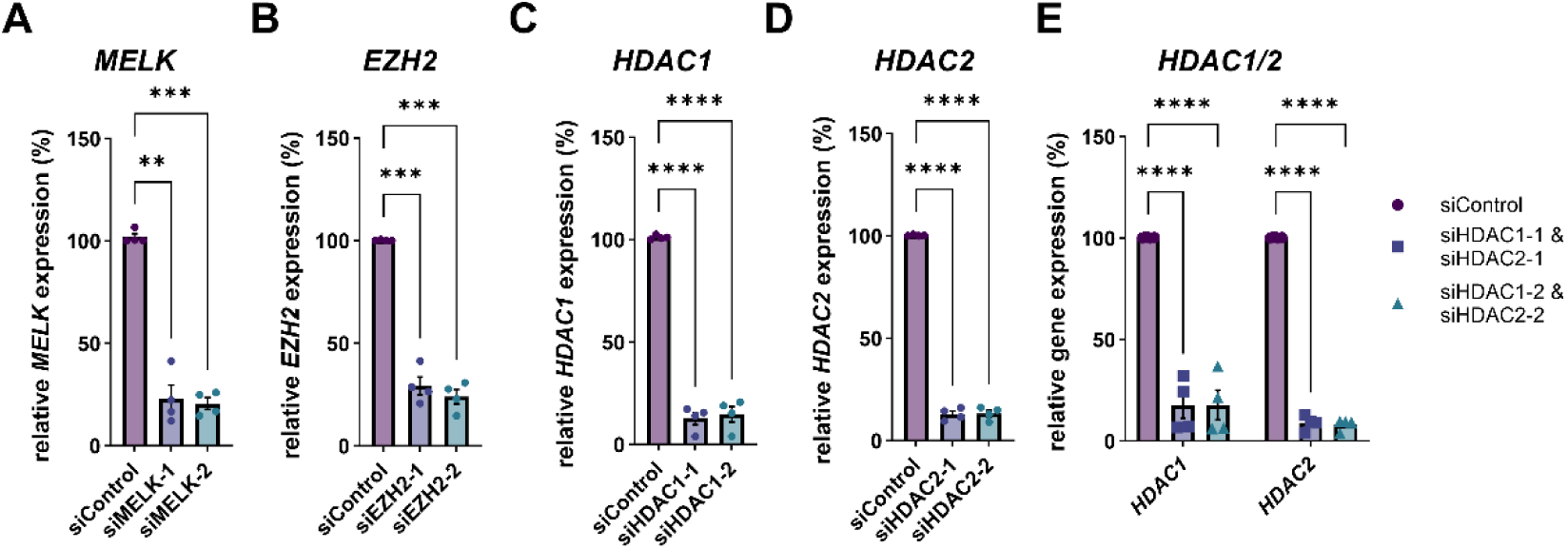
Gene expression upon knockdown of selected drug targets. RT-qPCR validation of RNAi-mediated knockdowns of **(A)** *MELK*, **(B)** *EZH2*, **(C)** *HDAC1,* **(D)** *HDAC2*, and **(E)** double knockout of *HDAC1/2* in four meningioma cell lines. *, *P* < 0.05; **, *P* < 0.01; ***, *P* < 0.001; ****, *P* < 0.0001.

**Suppl. Fig. 5:**
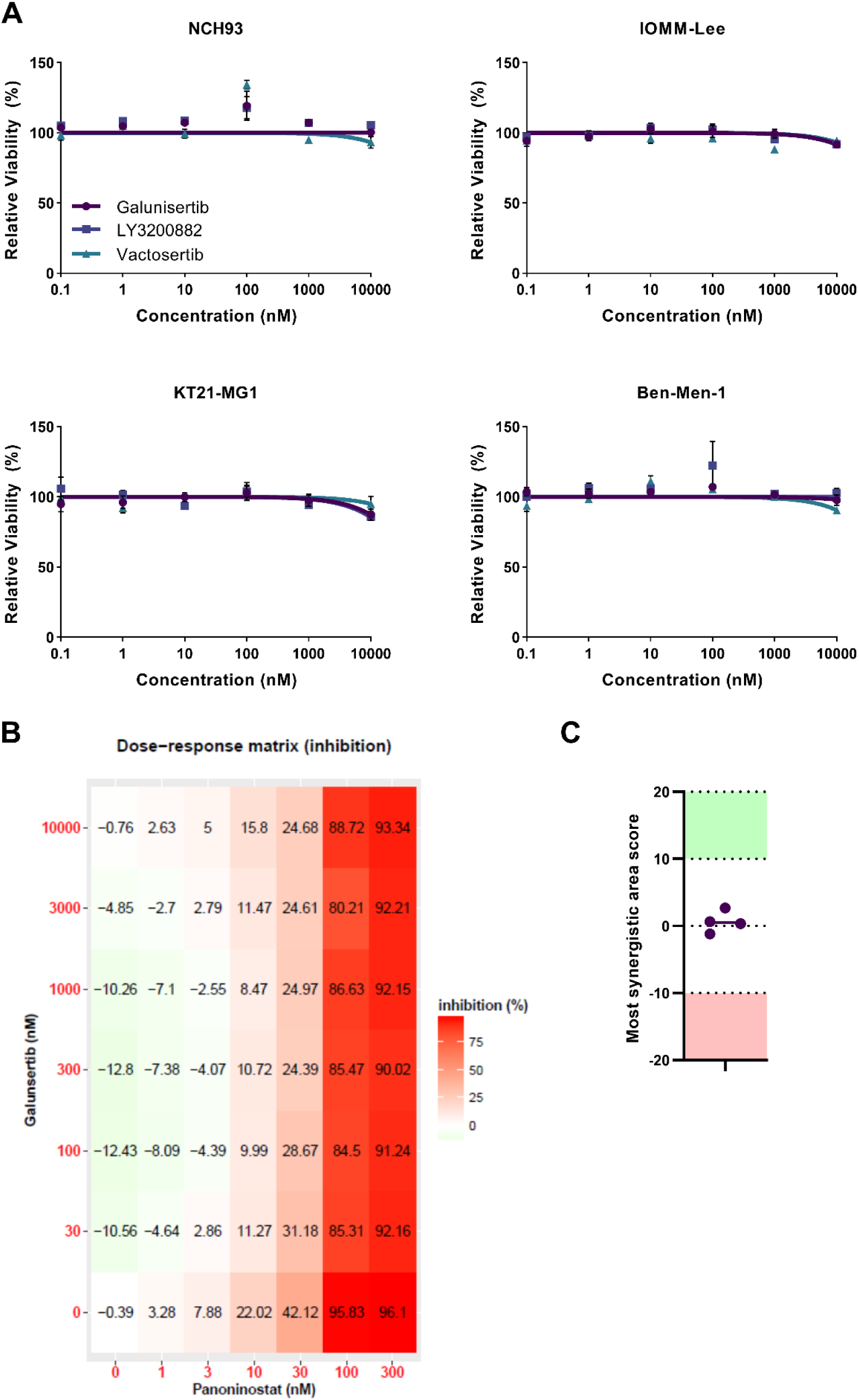
Treatment of meningioma cells with selected inhibitors. **(A)** Dose-response curves for TGFβi galunisertib, LY3200882 and vactosertib in four meningioma cell lines. IC_50_ values could not be established within the tested range, therefore single drug treatment had no impact on viability/proliferation of four meningioma cell lines. **(B)** Meningioma cells (NCH93) were treated in dose-response matrices with panobinostat from 1 to 300 nM and galunisertib from 30 to 10,000 nM. Synergy was calculated with Synergy Finder Plus in R. **(C)** Most synergistic area score from four meningioma cell lines. No synergy effect (synergy score >10) was observed when combining TGFβi galunisertib and HDACi panobinostat treatment.

## SUPPLEMENTARY TABLES

**Suppl. Table S1:** Library of targeted anticancer drugs.

| Name of drug | Target | Targeted Gene Name |
| --- | --- | --- |
| ABL001 | ABL1 | ABL Proto-Oncogene 1, Non-Receptor Tyrosine Kinase |
| Radotinib | ABL1 | ABL Proto-Oncogene 1, Non-Receptor Tyrosine Kinase |
| Brimonidine | ADRA2 | Alpha adrenergic receptor alpha-2 |
| StemRegenin 1 | AHR | Aryl hydrocarbon receptor |
| Perifosine | AKT | v-Akt murine thymoma viral oncogene homolog |
| Afuresertib | AKT1 | v-Akt murine thymoma viral oncogene homolog 1 |
| AT13148 | AKT1 | v-Akt murine thymoma viral oncogene homolog 1 |
| Miltefosine | AKT1 | v-Akt murine thymoma viral oncogene homolog 1 |
| Alectinib | ALK | Anaplastic lymphoma kinase |
| ASP3026 | ALK | Anaplastic lymphoma kinase |
| LDK378 | ALK | Anaplastic lymphoma kinase |
| AZD0156 | ATM | Ataxia telangiectasia mutated |
| ABT-263 | BCL2 | B-cell lymphoma 2 |
| AT-101 | BCL2 | B-cell lymphoma 2 |
| Bafetinib | BCR | Breakpoint cluster region protein |
| Rebastinib | BCR-ABL | Breakpoint cluster region-Abelson |
| Capivasertib | BRD2 | Bromo-2'-deoxyuridine |
| Acalabrutinib | BTK | Bruton's tyrosine kinase |
| Maraviroc | CCR5 | C-C chemokine receptor type 5 |
| Milciclib | CDK2 | Cyclin-dependent kinase 2 |
| Roscovitine | CDK2 | Cyclin-dependent kinase 2 |
| SNS-032 | CDK2 | Cyclin-dependent kinase 2 |
| Abemaciclib | CDK4 | Cyclin-dependent kinase 4 |
| Ribociclib | CDK4 | Cyclin-dependent kinase 4 |
| Palbociclib | CDK6 | Cyclin-dependent kinase 6 |
| Atuveciclib | CDK9 | Cyclin-dependent kinase 9 |
| Flavopiridol | CDK9 | Cyclin-dependent kinase 9 |
| MK-8776 | CHK1 | Checkpoint kinase 1 |
| Entacapone | COMT | Catechol-O-methyltransferase |
| Tolcapone | COMT | Catechol-O-methyltransferase |
| PF-477736 | CSF1R | Colony stimulating factor 1 receptor |
| PRN1371 | CSF1R | Colony stimulating factor 1 receptor |
| Reparixin | CXCR1 | C-X-C Motif Chemokine Receptor 1 |
| MSX-122 | CXCR4 | C-X-C Motif Chemokine Receptor 4 |
| Memantine | CYP2B6 | Cytochrome P450 Family 2 Subfamily B Member 6 |
| Itraconazole | CYP3A4 | Cytochrome P450 Family 3 Subfamily A Member 4 |
| Pioglitazone | CYP3A4 | Cytochrome P450 Family 3 Subfamily A Member 4 |
| Pinometostat | DOT1L | DOT1 like histone lysine methyltransferase |
| CPI-1205 | EZH1 | Enhancer of zeste homolog 1 |
| Tazemetostat | EZH2 | Enhancer of zeste homolog 2 |
| Derazantinib | FGFR | Fibroblast growth factor receptor |
| Debio1347 | FGFR2 | Fibroblast growth factor receptor 2 |
| JNJ-42756493 | FGFR2 | Fibroblast growth factor receptor 2 |
| LY2874455 | FGFR3 | Fibroblast growth factor receptor 3 |
| H3B-6527 | FGFR4 | Fibroblast growth factor receptor 4 |
| Linifanib | FLT/KDR | FMS-like tyrosine kinase/Kinase insert domain receptor |
| Dovitinib | FLT3 | FMS-like tyrosine kinase 3 |
| Pacritinib | FLT3 | FMS-like tyrosine kinase 3 |
| Tandutinib | FLT3 | FMS-like tyrosine kinase 3 |
| LY2090314 | GSK3A | Glycogen Synthase Kinase 3 Alpha |
| Entinostat | HDAC1 | Histone deacetylase 1 |
| Mocetinostat | HDAC1 | Histone deacetylase 1 |
| Panobinostat | HDAC1 | Histone deacetylase 1 |
| Resminostat | HDAC3 | Histone deacetylase 3 |
| Tasquinimod | HDAC4 | Histone deacetylase 4 |
| Ricolinostat | HDAC6 | Histone deacetylase 6 |
| 2-Methoxyestradiol | HIF1A | Hypoxia Inducible Factor 1 Subunit Alpha |
| BAY87-2243 | HIF1A | Hypoxia Inducible Factor 1 Subunit Alpha |
| Glycyrrhizic acid | HMGB1 | High Mobility Group Box 1 |
| BMS-986205 | IDO1 | Indoleamine 2,3-Dioxygenase 1 |
| Epacadostat | IDO1 | Indoleamine 2,3-Dioxygenase 1 |
| PF-06840003 | IDO1 | Indoleamine 2,3-Dioxygenase 1 |
| BMS-754807 | IGF1R | Insulin-like growth factor receptor |
| Linsitinib | IGF1R | Insulin-like growth factor receptor |
| Picropodophyllin | IGF1R | Insulin-like growth factor receptor |
| Itacitinib | JAK1 | Janus Kinase 1 |
| WP1066 | JAK2 | Janus Kinase 2 |
| AMG-510 | KRAS | KRAS Proto-Oncogene, GTPase |
| Lonafarnib | KRAS | KRAS Proto-Oncogene, GTPase |
| Minaprine | MAOA | Monoamine oxidase A deficiency |
| Pexmetinib | MAPK1 | Mitogen-activated protein kinase 1 |
| Ralimetinib | MAPK1 | Mitogen-activated protein kinase 1 |
| Idasanutlin | MDM2 | Mouse double minute 2 homolog |
| NVP-CGM097 | MDM2 | Mouse double minute 2 homolog |
| RG7112 | MDM2 | Mouse double minute 2 homolog |
| OTSSP167 | MELK | Maternal Embryonic Leucine Zipper Kinase |
| BMS-777607 | MET | MET Proto-Oncogene, Receptor Tyrosine Kinase |
| Foretinib | MET | MET Proto-Oncogene, Receptor Tyrosine Kinase |
| Tivantinib | MET | MET Proto-Oncogene, Receptor Tyrosine Kinase |
| Doxycycline | MMP1 | Matrix metalloproteinase-1 |
| Acetylcysteine | NFKB1 | Nuclear factor NF-kappa-B subunit 1 |
| KPT-9274 | PAK4 | P21 activated kinase 4 |
| Niraparib | PARP2 | Poly(ADP-Ribose) Polymerase 2 |
| Rucaparib | PARP2 | Poly(ADP-Ribose) Polymerase 2 |
| Veliparib | PARP2 | Poly(ADP-Ribose) Polymerase 2 |
| Pembrolizumab | PDCD1 | Programmed Cell Death 1 |
| Sildenafil Citrate | PDE5A | Phosphodiesterase 5A |
| Tadalafil | PDE5A | Phosphodiesterase 5A |
| Udenafil | PDE5A | Phosphodiesterase 5A |
| Crenolanib | PDGFRA | Platelet-derived growth factor receptor alpha |
| Masitinib | PDGFRA | Platelet-derived growth factor receptor alpha |
| Nintedanib | PDGFRA | Platelet-derived growth factor receptor alpha |
| AZD1208 | PIM1 | Pim-1 Proto-Oncogene, Serine/Threonine Kinase |
| INCB053914 | PIM1 | Pim-1 Proto-Oncogene, Serine/Threonine Kinase |
| BI 2536 | PLK1 | Polo Like Kinase 1 |
| Rigosertib | PLK1 | Polo Like Kinase 1 |
| Volasertib | PLK1 | Polo Like Kinase 1 |
| CFI-400945 | PLK4 | Polo Like Kinase 4 |
| GSK3326595 | PRMT5 | Protein arginine methyltransferases 5 |
| BMS-833923 | SMO | Smoothened/frizzled class receptor |
| Glasdegib | SMO | Smoothened/frizzled class receptor |
| Sonidegib | SMO | Spleen Associated Tyrosine Kinase |
| KX2-391 | SRC | SRC Proto-Oncogene, Non-Receptor Tyrosine Kinase |
| Saracatinib | SRC | SRC Proto-Oncogene, Non-Receptor Tyrosine Kinase |
| Entospletinib | SYK | Spleen Associated Tyrosine Kinase |
| Fostamatinib | SYK | Spleen Associated Tyrosine Kinase |
| TAK-659 | SYK | Spleen Associated Tyrosine Kinase |

**Suppl. Table S2:**
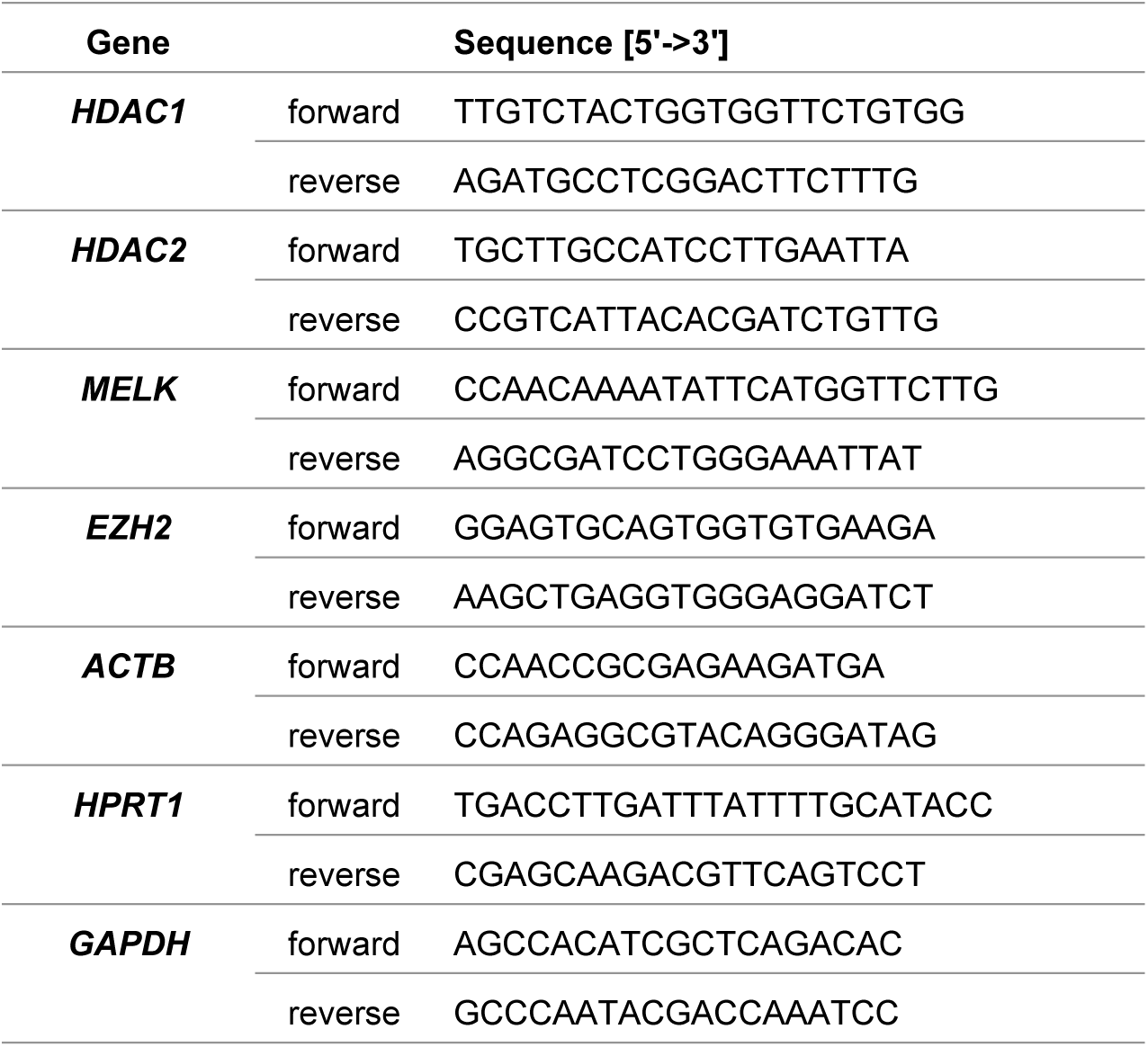
List of primers for qRT-PCR validation.

